# GRKs phosphorylate GPCR C-terminal peptides in a hierarchical manner

**DOI:** 10.1101/2025.05.06.652336

**Authors:** Arnelle Löbbert, Nils Lorz, Edda S. F. Matthees, Philip Rößler, Carsten Hoffmann, Alvar D. Gossert

## Abstract

Responses from G protein-coupled receptors (GPCRs) are downregulated in a precisely orchestrated process called desensitization. This process consists of two major steps: phosphorylation of the receptor by GPCR kinases (GRKs), predominantly on its C-terminus, and recruitment of arrestin, resulting in different signaling outcomes.

We carried out an NMR-based study of the phosphorylation patterns generated by GRK1 and GRK2 on C-terminal peptides of selected receptors (rhodopsin for GRK1, and β_1_- and β_2_-adrenergic receptors (ARs) for GRK2). Our data reveal that the kinases are promiscuous with respect to the substrate peptide, but produce clearly defined phosphorylation patterns on each substrate. We found pronounced differences in the rates at which certain residues are phosphorylated, in particular in the PXPP motifs in rhodopsin and β_1_AR. These results show, that GRKs produce well-defined phosphorylation patterns in absence of further modulators like the full receptor or Gβγ, and that the time profile of the phosphorylation barcode seems to be largely encoded in the minimal pair of C-terminal peptide and GRK. The data further suggest that arrestin might encounter different phosphorylation barcodes over time, potentially inducing different responses at different time points in the desensitization process.

## Introduction

G protein-coupled receptors (GPCRs) represent the largest family of transmembrane proteins and are targeted by approximately one third of all clinically employed drugs.(*1*) To control the signaling process of GPCRs, cells have developed a tightly regulated system of desensitization consisting of two major steps: (i) phosphorylation of the C-terminus and/or intracellular loops of the GPCR mainly through GPCR kinases (GRKs), and (ii) recruitment and binding of arrestins which in turn regulates a plethora of cellular responses.(*2*)

Interestingly, over 800 human GPCRs are known to be served by only seven GRK family members and four arrestins. This imbalance in diversity leads to the question of selectivity of GRKs as well as arrestins for distinct GPCRs. The barcode hypothesis, which states that phosphorylation of different sets of phosphorylation sites on the receptor triggers specific arrestin-mediated outcomes, has been proposed as a possible explanation.(*3*, *4*) In support of this hypothesis, two consensus findings have emerged regarding phosphorylation-dependent receptor-arrestin interactions. Firstly, a cluster of at least three close-by phosphorylation sites on the GPCR are required to change the conformation of arrestin to its receptor-activated state.(*5–8*) Patterns, such as PPP(*5*), PXPP(*7*, *8*), PXXPXXP(*6*) and PXPXXP(*6*), have been described to play a vital role in the binding of arrestin to receptor. Secondly, arrestin has been shown to dynamically employ distinct interaction interfaces to form complexes with GPCRs. It can either engage with the C-terminus of the receptor (“hanging conformation”) or it additionally binds to the intracellular cavity of the GPCR (“core conformation”), leading to distinct arrestin functions.(*9*, *10*) The latter complex conformation has been reported to be mainly triggered by receptor-proximal phosphorylation sites, while the hanging conformation predominantly occurs in combination with distal phosphorylation sites. At this point, it becomes crucial to understand the role of GRKs in producing such specific patterns. Although the identification of specific GRK-induced sets of phosphorylated sites has been the focus of extensive investigation for many years, the complex mechanism underlying GRK-catalyzed receptor phosphorylation remains unclear. What is firmly established, is that GRKs require activation by a receptor in its active state, that some kinases additionally require the G-protein βγ subunits for recruitment to the membrane, and that interactions with negatively charged phospholipids (phosphatidyl-inositol) enhance the phosphorylation reaction.(*11–15*) What is however debated, is whether the receptor domain or binding of Gβγ direct the GRKs to specific phosphorylation sites, or whether the final phosphorylation pattern is already encoded in the C-terminal amino acid sequence and the inherent specificity of the kinase.(*16*) Conversely this means, that GPCRs and other factors either stabilize the kinase in discrete states, or the receptor acts as an “on/off” switch of the kinase in a binary activation mechanism, where further factors could influence the reaction for example by enhanced recruitment of the kinase. It is also conceivable that the time-sequence of occurring phosphorylation events is relevant for the signaling outcome. However, experimental data to address these questions is difficult to obtain.

Early studies employing two-dimensional phosphopeptide mapping have provided initial insights into receptor phosphorylation in whole cells.(*3*, *17–19*) In the 2000s, the determination of individual amino acid phosphorylation of GPCRs by mass spectrometry (MS) became an attractive approach, enabled by rapid technical improvements. Thereby, numerous receptor phosphorylation sites have been identified with peptide or amino acid resolution.(*18–22*) Nowadays antibodies that recognize specific phosphorylated residues are commonly used to obtain new insights into the phosphorylation extent at the endpoint of a reaction, contributing to a quantitative understanding of receptor desensitization.(*23*) Still, phosphorylation data, and specifically time-resolved insights into the reaction, are relatively scarce: the most detailed studies have been carried out for bovine rhodopsin. Studies employing different methods by Ohguro, Zhang, and Kennedy *et al*.(*24–27*) showed that predominantly serine residues in the C-terminal tail are phosphorylated and that threonine residues are only phosphorylated to a lower extent. Azevedo *et al*. extended these finding by showing that threonine residues are phosphorylated more slowly, but are crucial for subsequent arrestin binding thereby indicating a precisely timed arrestin response.(*28*) For other receptor phosphorylation studies, maybe the study on the β2-adrenergic receptor by Nobles *et al*.(*4*) stands out. Here, a clear set of highly phosphorylated residues was identified, however without addressing the sequence of phosphorylation events which may be important to identify time-sensitive downstream responses.

In this study, we use NMR spectroscopy as it is a well-suited tool to investigate the chronological sequence of phosphorylation events with atomic detail. We employ a series of 2D NMR spectra to obtain time-resolved data on GPCR C-terminal phosphorylation. Using classical triple resonance experiments, we were able to assign the resonances to their respective amino acids and phospho-amino acids, such that site-specific data could be obtained. We follow the phosphorylation reaction of GRK1 and GRK2 on three different substrates: the C-terminal peptides of Rhodopsin (Rho Cterm), of β_1_ adrenergic receptor (β_1_AR Cterm) and β_2_ adrenergic receptor (β_2_AR Cterm). We chose these targets because they are widely studied, and because phosphorylation predominantly takes place on the C-terminus and not on intracellular loops of these receptors that are absent in our assay. Since the NMR experiments were performed *in vitro* with isolated kinases and peptides, we performed in-cell assays in parallel to confirm wild type-like activity of the used kinase constructs. With these validated tools at hand, we were able to observe the reaction on individual phosphorylation sites in a time-resolved manner, which allowed to reveal the sequence of phosphorylation events arising solely from kinase specificity and to identify individual dependencies in the phosphorylation hierarchy.

### Limitations of this study

Of course, also the use of NMR spectroscopy is associated with certain limitations. The presented study reveals the details of phosphorylation of isolated C-terminal peptides, but we were not able to directly compare the results to full-length receptors – despite attempting several approaches. Due to limited stability, the full-length systems are difficult to access for high-resolution kinetic *in vitro* studies. The current approach therefore only harnesses the basal activity of the enzyme. We use previously published results obtained in cells as a reference for the naturally produced phosphorylation patterns and compare it to the ones obtained *in vitro*. However, results derived from experiments performed in entire cells commonly do not dissect the contribution from individual kinases or other factors. Our results therefore provide clear data on the inherent specificity of GRKs. If cellular data identifies the same phosphorylation pattern, this suggests that the receptor only acts as a binary switch of the kinase. If the patterns deviate from literature, no conclusion can be drawn on whether altered specificities are caused by the receptor domain, other kinases, Gβγ or simply other alterations in the experimental setup.

## Results

### 1. Development of a time-resolved phosphorylation assay

In this study we established a GRK-dependent phosphorylation assay enabling time-resolved analysis of individual phosphorylation sites involved in GPCR desensitization (Figure 1). Peptides derived from the GPCR C-terminus of interest were used as surrogates for receptors, as the native full-length systems are hardly accessible for high-resolution kinetic *in vitro* studies due to their limited stability.

**Figure 1.**
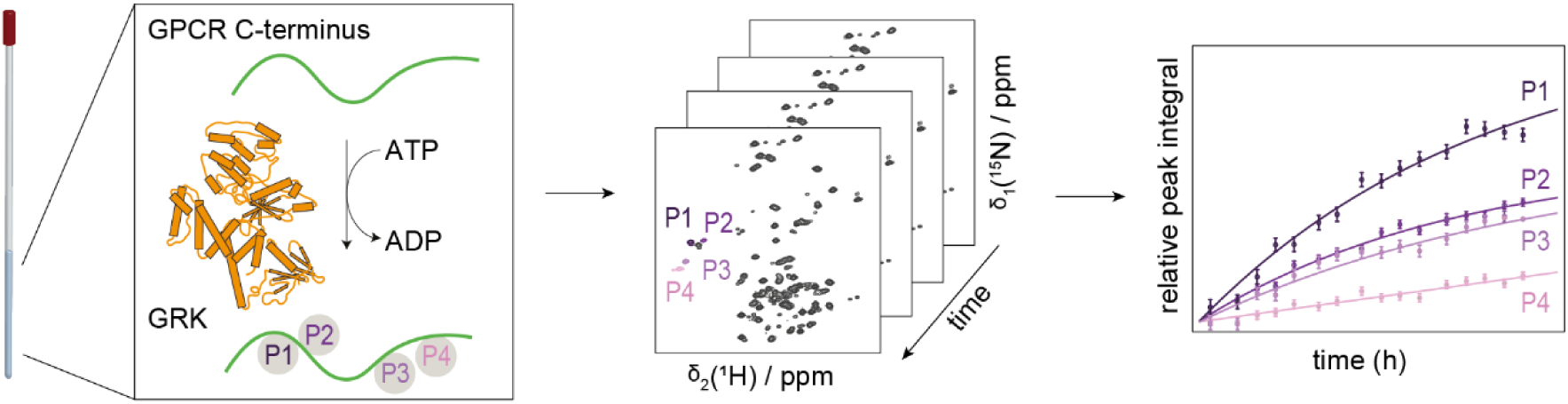
Schematic overview of the phosphorylation assay. To monitor the phosphorylation reaction by NMR spectroscopy, 100 µM ^15^N-labeled C-terminal peptide of the receptor of interest was mixed with 5 µM GRK and 10 mM ATP in a 3 mm NMR tube. A series of ^1^H-^15^N HMQC spectra, each recorded for 24 min, is subsequently measured every 4–6 h over the course of ∼60 h. The data is analyzed in Julia 1.9.2 to derive kinetic information on individual phosphorylation sites.

As additional negative charges induce measurable chemical shift perturbations in their surroundings, we employed NMR-spectroscopy to monitor the phosphorylation catalyzed by GRKs on ^15^N-labeled receptor C-termini. GPCR desensitization is a highly complex process, involving several effectors.(*29*) Here we aimed at establishing the basal phosphorylation, based solely on the GPCR Cterm-GRK interaction, in order to provide a reference for later studies on the influence of further effectors. Therefore, the reaction mixture only contained the compounds required to run this reaction in an NMR tube (100 µM substrate peptide, 5 µM GRK, 10 mM ATP, 25 mM MgCl_2_, 70 mM TRIS buffer, pH 7.2). A prerequisite for NMR research at atomic resolution is the assignment of resonances to their respective atomic nuclei. In contrast to large proteins, where resonance assignment is a major hurdle, peptide assignment can be easily achieved by classical three-dimensional NMR experiments. For all studied peptides, ≥90% of the backbone assignments in their unmodified form were obtained (Supplementary Table 2–4). Prolines were not included in this number, due to their lack of a backbone amide hydrogen, necessary for the assignment. In principle, measurement of consecutive 2D spectra enables a temporal resolution down to the minute time-scale. Since no additional factors were present in the reaction mixture, the reaction relied on the low basal activity level of the GRKs(*30*, *31*) resulting in a slow reaction rate. Hence, in our setup it was sufficient to measure 24 min 2D spectra at an interval of approximately 4 h, for 40–60 h.

To verify the ability of the GRK2 S670A construct employed in our assay to phosphorylate GPCRs under activated conditions, we compared the performance of GRK2 S670A to wildtype GRK2 (GRK2 WT) using two *in cellulo* assays, measuring Gα sequestration and arrestin recruitment. GRK2 has been shown to interact with the Gα_q_ and Gβγ subunits of activated G proteins.(*32*) This abolishes the interaction of G protein subunits with their effector molecules, such as phospholipase Cβ3 (PLCβ3) in case of Gα_q_. Using a split luciferase complementation assay (SLC)(*33*) in GRK2/3/5/6 knockout cells(*34*), we can assess the ability of the specific GRK construct to reduce the interaction between Gα_q_ and PLCβ3, which are fused to complimentary luciferase fragments. Here, both kinase variants (S670A and WT) showed equal Gα_q_ sequestration, reflected by the absence of a luminescence signal. In contrast, the control in GRK knockout cells transfected with the empty vector (EV) instead, Gα_q_-PLCβ interaction was measurable (Figure 2a–c). This results in an agonist concentration-dependent as well as time-dependent luminescence signal. The outcome of the SLC assay suggests correct folding and wildtype-like behavior of the used construct. However, this assay is independent of receptor phosphorylation. To ensure a WT-like outcome of receptor phosphorylation by GRK2 S670A, we performed a NanoBRET β-arrestin (βarr) recruitment assay in GRK knockout cells as elaborated in Drube *et al.*(*34*) (Figure 2e–f). Also here, the results display no significant alteration of the WT kinase compared to the used construct, neither in the concentration-response curves nor in the temporal profile. No βarr2 recruitment is obtained with EV, while both kinases lead to a highly similar extent of βarr2 recruitment. In-cell studies of the GRK2 S670A therefore confirm the full functionality of the used construct.

**Figure 2.**
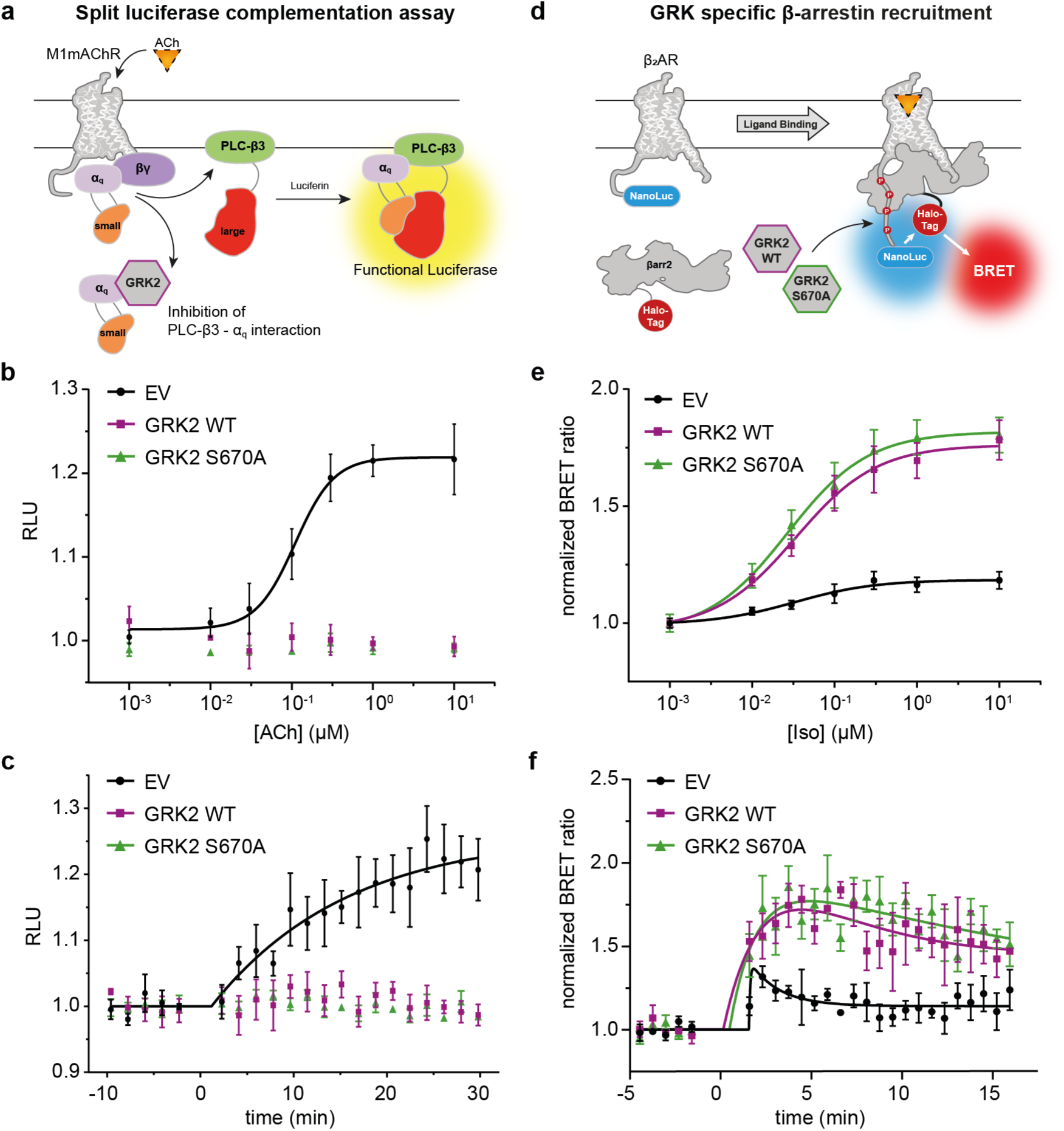
In cellulo assays confirm wild-type-like behavior of GRK2 S670A. (**a)** Schematic depiction of the split luciferase complementation assay (SLC).(33) Agonist activation of the Gα_q_-coupled M1mACh receptor leads to the dissociation of the G protein leading to Gα_q_-phospholipase C-β (PLCβ) interaction. The Gα_q_ subunit is tagged with the smaller fragment (orange) of the click-beetle luciferase whereas the larger fragment (red) is fused to PLC-β3. Using this split luciferase sensor, we monitored mAChR1-mediated Gα_q_-PLCβ interaction in ΔQ-GRK cells in the presence of GRK2 WT, GRK2 S670A and in absence of GRKs (empty vector (EV)-transfected). Data are analysed in ACh-concentration dependent (**b**) and time-dependent (**c**) manner as relative light units (RLU), mean of n = 4 independent experiments ± SEM. For concentration-dependent analysis data points were averaged over a time frame from 25-30 min (3 timepoints) post substrate addition. Time-dependent analysis is depicted for an ACh-concentration of 100 nM. **d** Schematic representation of the performed NanoBret β-arrestin (βarr) recruitment assay.(34) Agonist activation of the NanoLuciferase (NanoLuc)-tagged GPCR results in phosphorylation of the receptor by GRKs and subsequent recruitment of the Halo-Tag-βarr fusion protein. The recruitment-induced change in proximity of NanoLuc and Halo-Tag increases measured BRET ratios, enabling both, the agonist concentration-dependent as well as the time-dependent analysis of βarr recruitment. **e** GRK-dependent Halo-Tag-βarr2 recruitment to the β_2_AR-NanoLuc upon stimulation with isoprenaline (Iso) in ΔGRK2/3/5/6 HEK293 knockout cells (ΔQ-GRK). Cells were co-transfected with either EV, wild-type GRK2 (GRK2 WT) or GRK2 S670A. BRET data are presented as normalized BRET ratio, mean of n = 4 independent experiments ± SEM. **f** GRK-dependent Halo-Tag-βarr2 recruitment to the β2AR-NanoLuc shown over time at a stimulation level of 10 µM isoprenaline.

### 2. Investigating phosphorylation of Rho, β_1_AR, and β_2_AR C-termini by GRKs

The time-resolved NMR phosphorylation assay enables us to observe the phosphorylation on peptide substrates not only with temporal resolution, but also at atomic detail. We investigated in-depth the phosphorylation of the C-terminus of Rhodopsin (Rho) by GRK1 as well as the β_1_ adrenergic receptor (β_1_AR) and β_2_ adrenergic receptor (β_2_AR) by GRK2 (Figure 3) as these combinations represent the native pairs of C-terminus and kinase.

**Figure 3.**
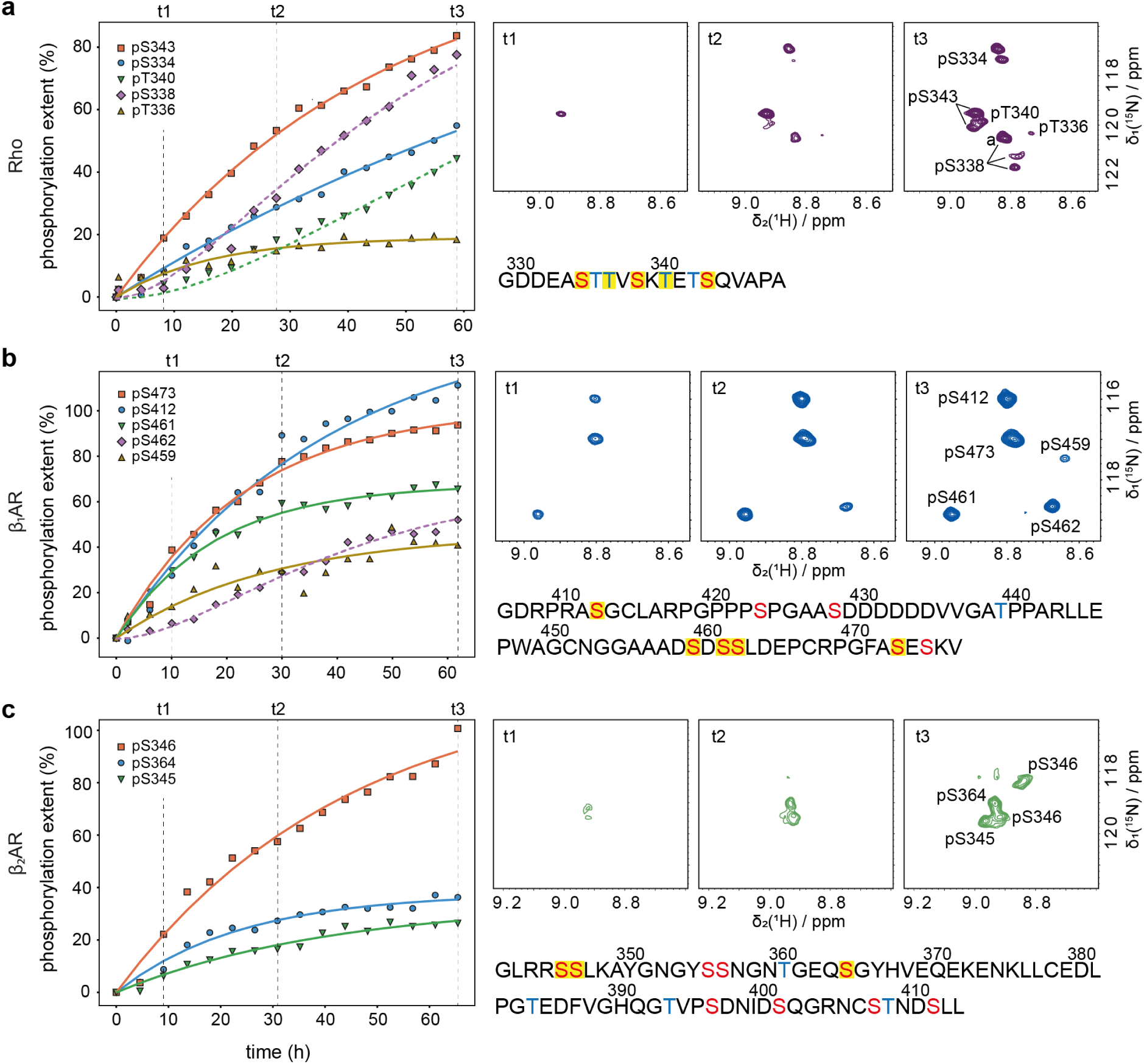
Phosphorylation of C-terminal peptides display different kinetics for individual phosphorylation sites. Phosphorylation reaction of Rho Cterm, catalyzed by GRK1 (**a**), β_1_AR Cterm, catalyzed by GRK2 (**b**), and β_2_AR Cterm, catalyzed by GRK2 (**c**). Peak integrals of individual sites are plotted over time (left). Peak integrals were normalized to their individual 100% phosphorylation level according to Theillet et al.(35) Data points were fitted to first-order kinetics (solid lines). Data which could not be fitted with classical first-order kinetics were subjected to analysis based on formulas 1.1–1.5 (Supplementary Figure 1). Fits are depicted with dashed lines (see figure 4 and supplementary figure 1 for details). Respective rate constants are summarized in supplementary table 5. On the top right of panels a–c, 2D [^15^N, ^1^H]-HMQC spectra displaying phosphorylated residues at individual timepoints are depicted. The last spectrum of each time-series displays the respective peak assignments. Sequence of C-terminal constructs are shown on the bottom right of panels a–c. Possible phosphorylation sites are marked in red (serines) or blue (threonines). Phosphorylation occurring in this assay are highlighted in yellow.

#### 2.1 Time-resolved analysis reveals phosphorylation hierarchy in each of the substrates

To analyse the results obtained in the NMR phosphorylation assay, peak integrals were plotted over reaction time. Hereby, individual kinetics of different phosphorylation sites became visible, enabling the determination of a chronological sequence of phosphorylation events.

Within the Rho Cterm, S343 is phosphorylated first. Then S334 gets phosphorylated, followed by S338 and T340 (Figure 3c). Interestingly, unlike the other phosphorylation curves in Rho Cterm, the data obtained for the specific phosphorylation of T340 and S338 cannot be fitted with classical first-order kinetics, but rather shows biphasic curvature, which we investigated in detail further below (Figure 4). Overall, the Rho Cterm is with its 20 amino acids the shortest tested C-terminus and offers 7 possible phosphorylation sites. In our assay, a total of 5 phosphorylated residues were observed. The Rho Cterm therefore exhibits a dense phosphorylation pattern and notably, it is the only substrate where strong threonine phosphorylation was observed.

**Figure 4.**
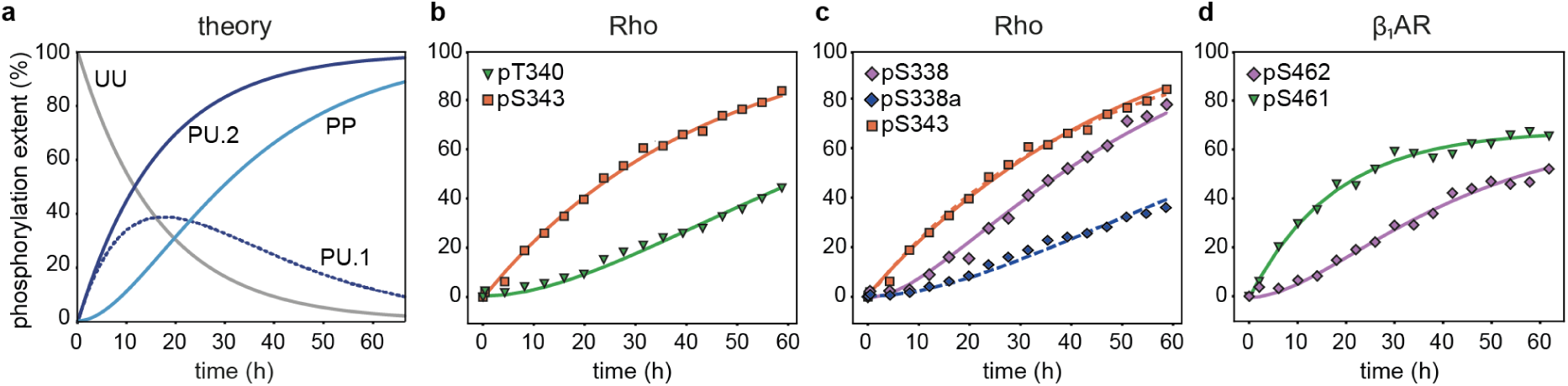
Fits based on a biphasic model indicate dependent phosphorylation events. (**a**) Simulation of normalized peak integrals based on formulas 1.1–1.5 (Supplementary Figure 1). Simulated curves are shown for different states, the unphosphorylated peptide (UU, gray), the mono-phosphorylated peptide (PU.1, dark blue, dashed line and PU.2, dark blue, solid line), and the doubly phosphorylated peptide (PP, light blue). PU.1 and PU.2 show the curve progressions for signals, for which the chemical shift is affected or unaffected by the second phosphorylation, respectively. For the simulation the following values were used: [UU_0_] =1, k_1_ = 0.06/h, k_2_ = 0.054/h. Fits are depicted for the phosphorylation pT340 in dependence of pS343 of Rho Cterm (**b**) and S462 in dependence of pS461 of the β_1_AR Cterm (**d**). For pS338, fits for the sum of all states (solid lines) as well as for pS338a only (dashed lines) are shown in dependence of pS343 of the Rho Cterm (**c**) (see text for details).

For the β_1_AR Cterm, the first phosphorylated site was S473, followed by S412, and afterwards S461. The last two sites phosphorylated by GRK2 were S462 and S459 (Figure 3a). Similar to T340 and S338 of the Rho Cterm, the phosphorylation of S462 cannot be fitted by first-order kinetics. In total, we observed that 5 of 9 serine residues in β_1_AR Cterm were preferentially phosphorylated by GRK2 whereas threonine phosphorylation was not observed at all. Further, these phosphorylations occurred at different rates, including a peculiar non-first-order phosphorylation behavior of S462.

The most prominent phosphorylation sites in β_2_AR Cterm are S364 and S346, followed by S345 (Figure 3b). All curves could be fitted based on first-order kinetics. Remarkably, clearly fewer sites get phosphorylated than in β_1_AR Cterm, even though the β_2_AR Cterm offers a larger number of possible phosphorylation sites (13 vs. 9). Furthermore, in β_1_AR Cterm, phosphorylations happen rather evenly distributed over the C-terminal tail, but in β_2_AR Cterm only sites that would be rather proximal to the receptor are phosphorylated (Figure 3b).

#### 2.2 A biphasic model as alternative fitting method indicates dependent phosphorylation events

Several phosphorylation curves could not be fitted with first-order kinetics, for instance T340 and S338 in Rho Cterm as well as S462 in β_1_AR Cterm. To investigate the biphasic curvature of these phosphorylation curves, we explored a model for dependent reactions, where the existence of a first phosphorylation site is a prerequisite for the phosphorylation of the second site (Supplementary Figure 1, Eq. 1.1–1.5). In figure 4a, simulated curves are shown based on the described model. Two different options are possible for the monophosphorylated state, termed PU (for one phosphorylated, P, and one unphosphorylated residue, U). In the first scenario, the phosphorylation of the second site leads to chemical shift perturbation of PU and therefore in a decrease of the peak integral of PU (curve PU.1 in figure 4a). In the second case, the second phosphorylation site does not affect the chemical shift of PU, therefore peaks keep increasing until saturation (curve PU.2 in figure 4a).

To identify the dependency of the phosphorylation of a certain site on another phosphorylation site all present peaks were subjected to analysis based on the described model. For T340 in Rho Cterm, using the RMSD as an evaluation parameter, the best fit was achieved in combination with the data obtained for pS343 (Figure 4b). Similarly, S462 in β_1_AR Cterm seems to be triggered by prior phosphorylation of S343 (Figure 4d). It is however, important to mention that the combination with several first site candidates yielded reasonable fits and we lack a clear cut-off criterium (see supplementary figure 3 for variability of the data). For instance, in Rho Cterm S343 and S334 gave similar results as triggers for T340, and in β_1_AR Cterm S473 gave worse but still acceptable fits in combination with S462. However, in an unfolded peptide it is more likely that amino acids in close sequential proximity have an impact, therefore, in figure 4b–d the best fits with closest neighbors are shown.

At a first glance, the phosphorylation of S338 in the Rho Cterm seems also to be dependent on the phosphorylation of S343. However, several states of pS338 exist (Figure 3a, t3) and interestingly, the rise of pS338a is highly similar to the one of pT340. This indicates that S338 is phosphorylated independently of pS343, but the phosphorylation of T340 results in a peak shift of pS338, which imprints its non-monoexponential kinetics on pS338a.

In summary, we identified three phosphorylation sites, which could not be fitted with first-order kinetics. Our attempt to fit a biphasic model suggest dependencies of S462 phosphorylation on S461 in β_1_AR, and T340 phosphorylation on S343 in Rho.

#### 2.3 Phosphorylation of different lengths of C-terminal peptides yield consistent results

Our NMR assay in this simplified system allowed us to identify time-resolved phosphorylation of isolated C-terminal peptides by GRKs. However, many aspects may not represent the situation *in vivo*. For instance, in a peptidic context it might be that just the most N- or C-terminal residue is preferentially phosphorylated. To assess such potential artefacts, we tested different lengths of the β_2_AR C-terminal peptide, which contains two SS motives (S345/S346 and S355/S356) (Supplementary Figure 4). Constructs with a four amino acid N-terminal overhang prior to the respective SS motive revealed near exclusive phosphorylation of the S345/ S346 motif. S355 and S356 were not phosphorylated regardless of their position in the peptide, while for example S364 was consistently phosphorylated in all three different constructs. These results therefore indicate a strong sequence specificity and little, if any, influence of the peptide length.

#### 2.4 Comparison of GRK1 and GRK2 phosphorylation

In order to obtain a more detailed understanding of the phosphorylation behavior of GRK1 and GRK2, we tested the reaction of both kinases on all three substrates (Figure 5). For comparison of the phosphorylation driven by the two isoforms, we analyzed the relative phosphorylation extent on individual sites of the different C-termini at the endpoint of the reaction (after 60 h).

**Figure 5.**
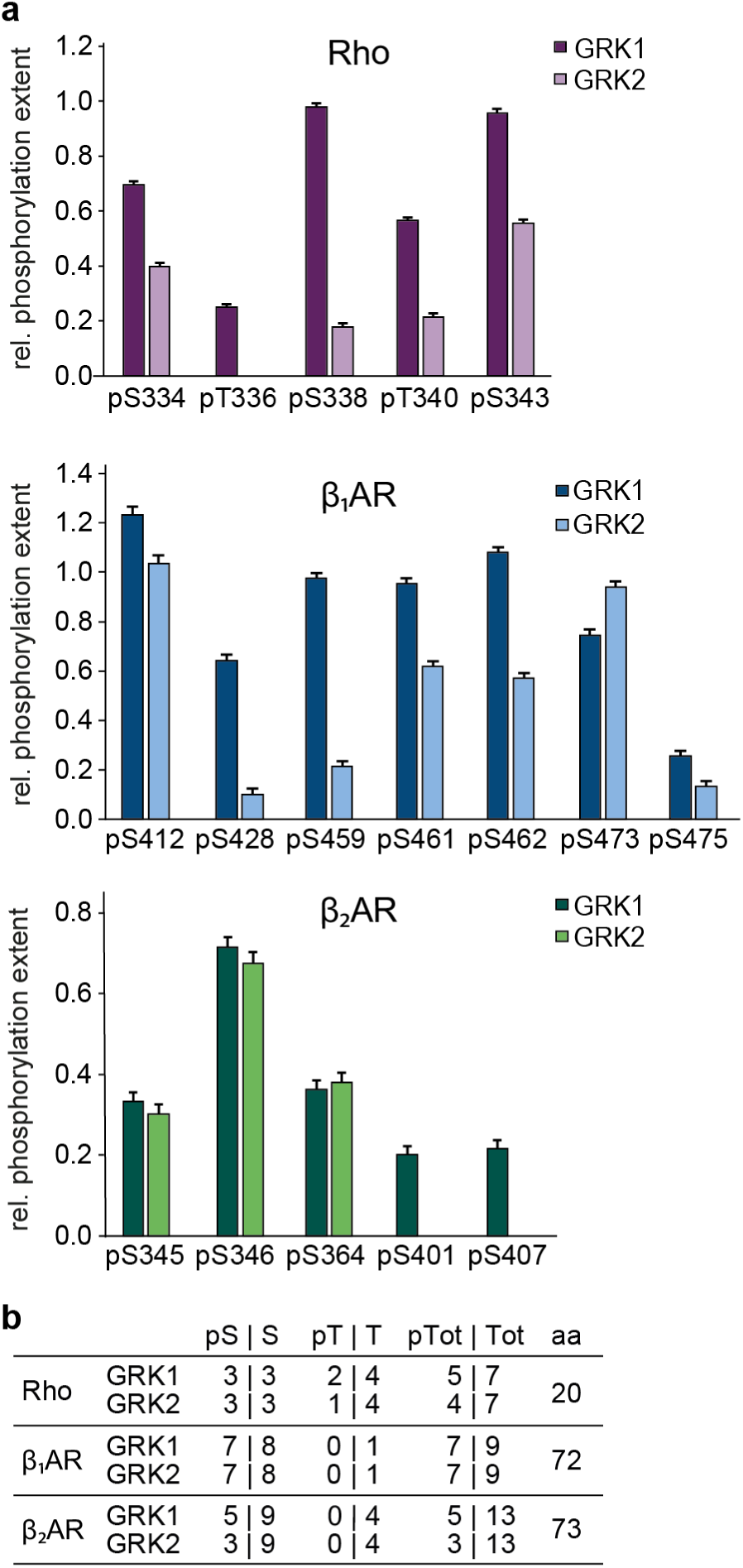
Comparison of GRK1- and GRK2-catalysed reactions reveal higher phosphorylation degree with GRK1 and promiscuity of both kinases. (**a**) Bar graphs depicting the phosphorylation of Rho (purple), β_1_AR (blue), and β_2_AR (green) by either GRK1 (darker tone) or GRK2 (lighter tone). Endpoints of each reaction (at ∼60 h) were chosen for analysis. Error bars indicate the standard deviation of noise of the respective spectra. For all three peptides, GRK1 shows enhanced phosphorylation activity compared to GRK2. (**b**) Phosphorylation sites of each peptide. pS = phosphorylated serines by GRK1/2, pT = phosphorylated threonines by GRK1/2, pTot = total phosphorylated residues, aa = total number of amino acids. Both kinases prefer serine residues over threonine residues.

Phosphorylation events were visible in all six assays, indicating that both GRKs are not specific to their native targets (Figure 5a). Notably, GRK1 and GRK2 displayed highly similar site selectivity, but GRK1 is able to phosphorylate the given substrates to a higher degree than GRK2. For the Rho Cterm as well as for β_2_AR Cterm, GRK1 is able to phosphorylate additional residues, which are not phosphorylated by GRK2. There are different possible ways to interpret these results: Either GRK1 shows less selectivity than GRK2 or the faster reaction rate of GRK1 compared to GRK2 leads to the appearance of phosphorylated residues which cannot be seen within our timeframe for GRK2-driven reactions.

Further, our data show that serine residues are preferred over threonine residues by both GRKs (Figure 5b). Within our assay threonine residues were only phosphorylated in the Rhodopsin C-terminus. The comparison of the phosphorylation of all three C-termini by both GRKs further reveals, that in the β_2_AR Cterm the least sites were phosphorylated compared to β_1_AR and Rho Cterm (Figure 5b). The different number of phosphorylated sites of β_1_AR and β_2_AR Cterm is especially interesting as the constructs are of similar length and the β_1_AR Cterm has only 9 potential phosphorylation sites, whereas the β_2_AR Cterm offers 13 options for phosphorylation. This, in combination with the results shown in figure 3, where it becomes evident, that the β_1_AR Cterm is phosphorylated to a higher degree compared to the β_2_AR Cterm by GRK2, leads us to the conclusion that the β_1_AR Cterm seems to be a better substrate for the kinase.

## Discussion

The establishment of an NMR-based time-resolved phosphorylation assay with residue resolution enabled the investigation of the phosphorylation pattern of three different GPCR C-termini. The *in vitro* study of the C-terminal stretches of the Rho C-terminus, which is naturally phosphorylated by GRK1, as well as β_1_AR and β_2_AR, which are native substrates of the GRK2 isoform, revealed distinct phosphorylation patterns, and more intriguingly, a phosphorylation hierarchy in each of the different C-termini. This provided valuable insights into the functioning of GRKs.

Comparing the phosphorylation of the three substrates by either GRK1 or GRK2 revealed that both kinases are not specific to their native peptide substrates (Figure 5). This highlights the importance of tissue- or cell-specific expression patterns for each isoform.(*36*) Furthermore, the two kinases yield similar phosphorylation patterns, but GRK1 was shown to be more prolific on the individual substrates. The presented assay reports on the basal activities of the individual kinases as we focused on C-terminal receptor peptides rather than stimulated receptors that would activate the GRKs. The results therefore indicate that GRK2 has a lower basal activity compared to GRK1 (Figure 5). The different basal activities could be attributed to the fact that GRK1 is the predominantly expressed isoform in retinal rod cells(*37*), whereas GRK2 is expressed ubiquitously in most other cells of the body alongside GRK3, GRK5 and GRK6.(*36*, *37*) The coexistence of four isoforms in one cell most likely requires stronger regulation of the different kinases compared to GRK1, resulting in a lower basal activity of GRK2. Furthermore, GRK1 regulates the extremely fast visual process, which requires a prolific enzyme, compared to most other receptor responses, which can be regulated by other GRKs at a more moderate pace. Interestingly, in contrast to GRK2, GRK1 undergoes significant autophosphorylation at several sites (S5, T8, S21, S488, T489) in the presence of ATP.(*38*) However, it is suggested, that the autophosphorylation has no major impact on the phosphorylation reaction on soluble substrates.(*38*, *39*)

Even though the GRKs are promiscuous, they do display site-selectivity and yield reproducible phosphorylation patterns on each C-terminus (Figure 6a). In our assay, GRK2 displays different phosphorylation activity on both its native substrates. While in the β_1_AR Cterm 5 of 9 possible sites were phosphorylated, the β_2_AR Cterm was phosphorylated only on 3 sites, although 13 Ser and Thr residues are present. A possible explanation for this could be the negative patch in the sequence of the β_1_AR (Figure 3a), which is beneficial with regard to the acidophilic nature of the kinase.(*40*, *41*) Therefore, the stretch of six aspartates could enhance the affinity of the kinase towards the β_1_AR C-terminal tail, compared to the β_2_AR C-terminus. Further, we see a strong preference of both kinases for serine over threonine residues which goes in line with the results obtained by Onorato *et al.*(*40*) as well as Johnson *et al.*(*41*) This emphasizes the crucial role of the amino acid sequence of the substrate in the phosphorylation activity of the GRKs even/already without the impact of a receptor core.

**Figure 6.**
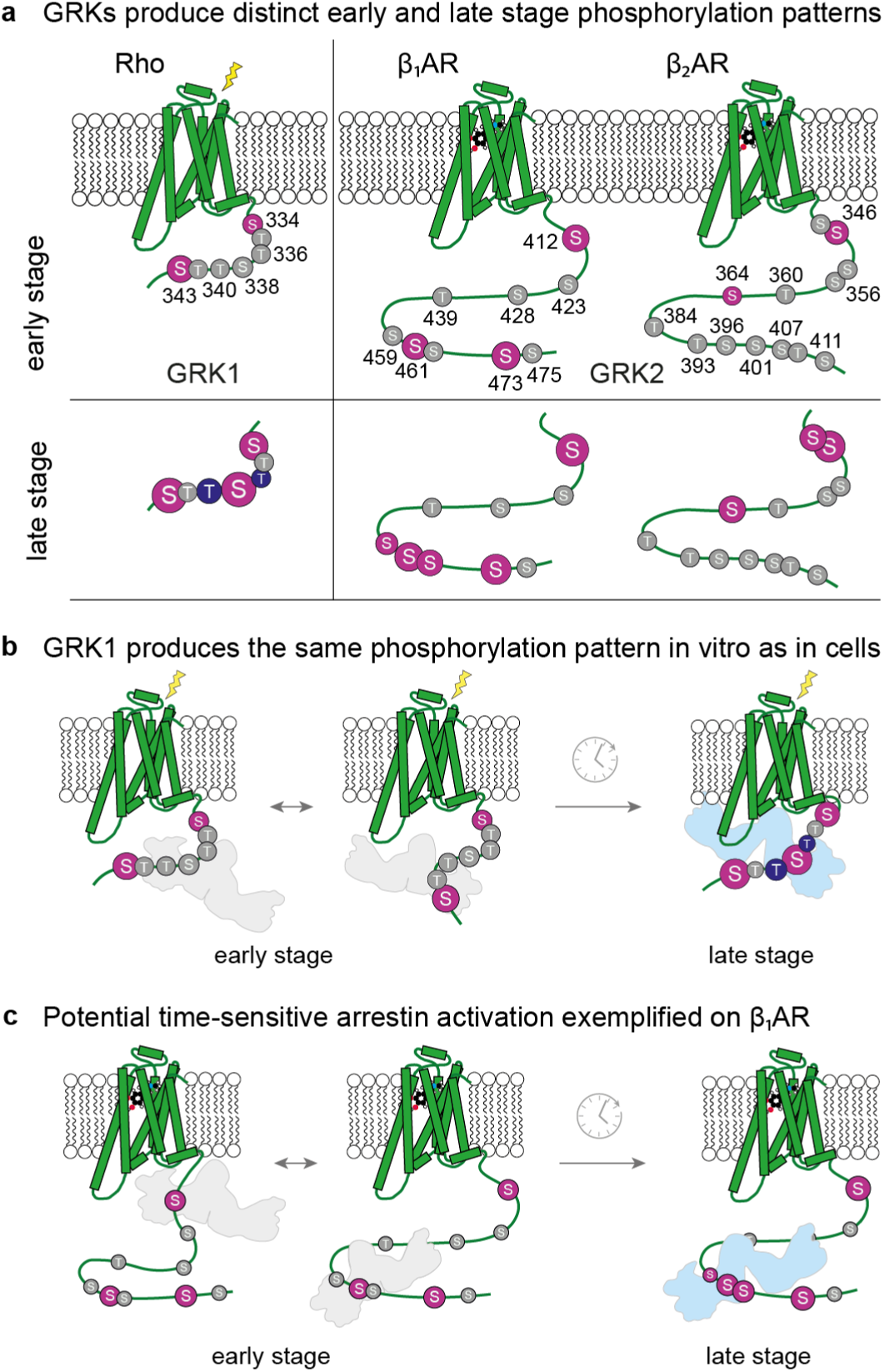
Schematic representation of distinct early and late-stage C-terminal phosphorylation by GRK1 and GRK2. (**a**) Results obtained at an early stage (∼10 h) vs a later stage (∼60 h) in the phosphorylation assay were indicated by color and size on the respective C-terminus. Possible phosphorylation sites are marked in gray. Phosphorylated serine residues are highlighted in pink and phosphorylated threonine residues in blue. The size of the respective sites indicates the degree of phosphorylation. (**b, c**) Possible interaction modes of arrestin (gray = inactive, light blue = active) with the different phosphorylation states of a given C-terminus are shown. (**b**) The phosphorylation hierarchy of the Rho C-terminus observed in cells is maintained in the in-vitro assay. Slower phosphorylation of the threonine residues which are crucial for arrestin interaction lead to a delayed arrestin response. (**c**) Proposed time-sensitive arrestin activation is shown representatively for the β_1_AR. Here, the delayed phosphorylation of the arrestin-activating PXPP motif may act as a timer for specific arrestin responses.

Taking a closer look at the individual phosphorylation sites indicates a switch-like (on/off) activation mechanism of the kinase by GPCRs, rather than the receptor shaping the kinase specificity. The phosphorylation of rhodopsin is a well-studied system. In rod cells, only one GRK family member is present, namely GRK1. This natural isolation of the receptor-kinase pair renders the system highly suitable for an *in vitro* replication as well as sensible comparison of the results to in-cell data. We therefore selected this system to assess the transferability of the results obtained on peptide substrates to full-length receptors. In our assay, Rho Cterm is predominantly phosphorylated on its three serine residues S343, S338 and S334, followed by T340 and T336. This goes in line with results obtained on full-length receptors (Supplementary Figure 5). *In vivo* studies from Ohguro *et al*. demonstrated phosphorylation of S334, S338, T336 and S343.(*25*) Here, S334 and S338 were found to be the main sites of phosphorylation, both of which are readily phosphorylated in our assay (Figure 3). Studies on the purified full-length receptor by Zhang *et al*. describe the phosphorylation of S334/T335/T336 (individual sites not dissected), S338, T340 and S343, again supporting the results obtained in our assay.(*26*) The studies from Kennedy *et al.* and Azevedo *et al.*, showed in their studies predominant phosphorylation on the three serine residues.(*27*, *28*) Both studies further suggest slower phosphorylation of threonine residues. Intriguingly, our results indicate a delayed phosphorylation behavior of T340 and show that T336 is phosphorylated only to a small extent (Figure 4). It is remarkable that our simplified assay on C-terminal peptides results in highly similar phosphorylation outcomes compared to the phosphorylation of full-length receptors. The high resemblance of the phosphorylated sites combined with the retained fast and slow phosphorylation of serine and threonine residues, respectively, indicate that the kinase may be activated in a switch-like manner by the receptor, rather than the receptor influencing the kinase specificity.

To further test this hypothesis, we turned to a literature comparison of the observed GRK2 phosphorylation patterns on its native substrates, the β_1_AR Cterm and β_2_AR Cterm. However, this represents a more complex scenario. In most human cells, GRK2, GRK3, GRK5 and GRK6 are co-involved in the process of receptor phosphorylation and time-resolved data on distinct phosphorylation events catalyzed by individual kinases is currently lacking. Hence the comparability of the obtained results with in-cell studies is limited and only obtained endpoints could be compared.

The investigation by Hinz *et al.* of the β_1_AR (seemingly the only current phosphorylation analysis on this receptor), revealed the phosphorylation of S412 and S461/S462 which is consistent with our assay.(*42*) However, we did not observe phosphorylation of S423 which is reported in the study. As the analysis of phosphorylation events was analyzed on a purified receptor from cells, the contribution of the individual kinases could not be differentiated. The discrepancy of some phosphorylation sites might therefore arise from different kinase contributions or the specificity of GRK2 may be altered by the receptor or other effectors in the cell. However, in light of the obtained results on the phosphorylation of Rho Cterm by GRK1, we consider the first scenario more likely.

Trester-Zedlitz *et al.* analyzed the phosphorylation pattern on β_2_AR derived from overexpressing HEK293 cells after agonist-stimulation and showed that the β_2_AR Cterm is not phosphorylated on distal sites, which is consistent with our results.(*20*) In contrast, Shiraishi *et al*. as well as Nobles *et al*. demonstrated the phosphorylation of two rather distal sites of the β_2_AR, S401 and S407, by GRK2.(*4*, *43*) In our assay, which is limited by the stability of the kinases over the long reaction time, these sites were not phosphorylated above the detection threshold of roughly 5% by GRK2. Given that β_2_AR Cterm seems to be a poor substrate for GRK2 (as described before), combined with the low basal activity of the GRK family member, these sites could potentially also appear after longer reaction times or activation by stimulated receptors.

In general, the comparison of our end-point results to studies including *in vitro* phosphorylation of full-length receptors and in-cell assays analyzed by mass spectrometry or phosphosite-specific antibodies reveals considerable similarity between the obtained data, supporting the hypothesis of a switch-like activation model for the kinase by the receptor. This in turn, increases the confidence that the hierarchical phosphorylation events observed in our assay may represent a physiologically relevant scenario, with the exception of overall rates of the reaction. Through activation of the GRKs by interaction with an activated receptor, the reaction rates will be drastically increased, while the specificity and therefore the chronological sequence of receptor phosphorylation may be maintained.

The time resolution of our assay thus provides valuable insights into GRK function. A first simple observation is that the different phosphorylation sites display strongly different rates. This clearly shows, that an individual kinase does not bind once to the peptide and slides on it thereby phosphorylating all potential sites in one event before dissociating again. The kinase rather binds and dissociates several times from the peptide with different likelihoods of phosphorylating different residues. This produces chronologically different phosphorylation patterns.

Considering the observed chronological phosphorylation events from the perspective of downstream signaling, this opens up the possibility of time-sensitive arrestin responses. Studies on arrestin– rhodopsin interaction revealed that rapid serine phosphorylation on rhodopsin does not lead to full deactivation of the receptor.(*28*) Only the slower phosphorylation of threonine residues effectively promotes arrestin binding, fully deactivating the receptor (Figure 6b). It is intriguing that in our simplified assay, T340 displays such a delayed phosphorylation behavior. Furthermore, even though the Rhodopsin C-terminus is strongly phosphorylated compared to the other two substrates within our assay, T342 is not phosphorylated at all. Interestingly, this site is postulated as “inhibitory site” by Mayer *et al.*(*44*) In line with this hypothesis, it is sensible that GRK1 does not target T342 at an early stage. Together, the results suggest a tight control of the timing of rhodopsin desensitization. In our study, we extend this hypothesis beyond the nature of the phosphorylated residue type (Ser/Thr) by the specificity of the kinase, which entails delayed phosphorylation of individual residues.

By taking a closer look at the phosphorylation pattern of the β_1_AR Cterm, it becomes apparent that first three sites that are quite widely dispersed are phosphorylated by GRK2 (S473, S412, then S461). As the activation of arrestin requires the phosphorylation of three sites which are close in space(*5–7*), the protein will not be activated at this stage of phosphorylation (Figure 6c). At a later time-point – and most notably in likely dependence of S461 – S462 is phosphorylated. Also, S459 is phosphorylated to a small extent. These three serines form a PXPP motif, which has recently been described as an important binding and activation motif for arrestin.(*7*, *8*) It is noteworthy that the kinase does not initially target all serine residues in the PXPP motif and that the phosphorylation of S462 depends on the adjacent phosphorylation site. This could hint at a time-delay in the arrestin response which is proposed to be triggered by the PXPP motif. It is intriguing to speculate at this point that arrestin may show an early and a late stage response in dependence to the different prevailing phosphorylation patterns, which change over time (Figure 6). A previous study proposed that agonist-dependent stepwise phosphorylation of the leukotriene B_4_ type-1 receptor yields distinct arrestin complexes, leading to diverse cellular functions.(*45*) Also, Haider *et al*., demonstrated in their *in cellulo* work that arrestins can adopt different conformations arising from distinct phosphorylation patterns on the same GPCR.(*46*) Both studies confirm that a single receptor can induce different arrestin states based on varying phosphorylation patterns.

Recently it has been reported, that arrestin can engage with the receptor in two different manners which are dependent on the location of the phosphorylation sites.(*9*, *10*) It can either adopt a hanging conformation, which is proposed to be facilitated by distal phosphorylation sites, or a core conformation, which is promoted by proximal phosphorylation sites. Our data suggest that by the end of the reaction the β_1_AR Cterm is phosphorylated rather on distal sites (Figure 6a). In contrast, the β_2_AR C-terminal domain is mainly phosphorylated on proximal sites, even though a range of distal phosphorylation sites are present. This could indicate different arrestin binding modes for the two receptors at a given time point.

In conclusion, our study highlights the intricacy of the GRK-dependent phosphorylation reaction in the desensitization process of GPCRs. We could show that the kinases are non-specific to C-terminal peptides, but display site-selective phosphorylation of those. The amino acid sequence of the substrates significantly impacts the activity of the kinase and governs the site-selectivity of the enzymes beyond the mere distribution of serine and threonine residues as shown for the β_1_AR Cterm in comparison to the β_2_AR Cterm. The comparison of our simplified assay with literature data on full-length receptors indicates a switch-like activation mechanism for the kinase by the receptor. Conversely, this strengthens the confidence that the presented results may reflect a physiologically relevant scenario. Our assay shows that arrestin binding motives, such as the PXPP motif, may be present but not necessarily phosphorylated at an early stage. The chronological sequence of the phosphorylation events thereby adds another layer of complexity to the GPCR inactivation process, opening up the possibility for time-delayed responses and for the transition from one arrestin state to another over time. Hence, understanding the GRK-perspective of desensitization is crucial to unravel the mystery of the inactivation of hundreds of GPCRs by just a handful of proteins.

## Material and Methods

### 1. Expression and purification of ^15^N and ^15^N,^13^C labeled C-termini

Plasmid DNA coding for the C-termini of *bt*Rhodopsin (329–348), *hs*β_1_AR (406–477), *hs*β_2_AR01/02/03 (342–413/351–413/358–413) (pET28b-based) were transformed into *E. coli* BL21(DE3) Codon Plus RIL cells (Agilent). Cells were grown on an agar plate supplemented with chloramphenicol and kanamycin at 37 °C overnight. A colony was picked to inoculate 2 mL of LB medium. Cells were incubated in a shaker at 37 °C for 4 h at 180 rpm. The mixture was then transferred into 50 mL of M9 minimal medium, containing either 1 g/l ^15^NH_4_Cl and 4 g/l glucose or 1 g/l ^15^NH_4_Cl and 3 g/l glucose-^13^C_6_ for U-[^15^N] or U-[^13^C,^15^N] labeled protein, respectively. The preculture was incubated overnight at 37 °C and 180 rpm. For the main culture, 950 mL of the previously mentioned medium was prewarmed to 37 °C and the preculture was added. The culture was shaken and grown at 37 °C until the OD_600_ reached 0.6 at which point the temperature of the shaker was set to 20 °C. Half an hour later, 1 mM isopropyl β-D-1-thiogalactopyranoside (IPTG) was added. Cells were harvested the next day by centrifugation for 10 min at 6,000 × g at 4 °C.

For purification, cells were lysed in lysis buffer (50 mM TRIS pH 8.0, 300 mM NaCl, 5 mM imidazole, 1 mM PMSF, DNaseI and 10% v/v glycerol) with an LM10 Microfluidizer (Microfluidics) and the obtained lysate was centrifuged for 1 h at 35,000 × g at 4 °C. The supernatant was filtered with a 0.45 µm syringe filter (Sarstedt) and loaded onto a 5 mL Ni-NTA Superflow-Cartridge (Qiagen). The column was washed with washing buffer (50 mM TRIS pH 8.0, 300 mM NaCl, 5 mM imidazole, 10% v/v glycerol, 1mM DTT) and the protein was step eluted with elution buffer (50 mM TRIS pH 8.0, 300 mM NaCl, 250 mM imidazole and 10% v/v glycerol, 1mM DTT). A buffer exchange to washing buffer was conducted using a HiPrep 26/10 Desalting Column (GE). TEV protease was added and after overnight incubation a reverse Ni-IMAC purification step was performed to remove TEV protease and cleaved tag. The peptide solutions were loaded onto a Superdex 75 Increase 10/300 column (GE) to exchange the buffer to 50 mM TRIS pH 7.2, 150 mM NaCl, 25 mM MgCl_2_. The peptide containing fractions were concentrated in 3 kDa Amicon-Ultra 4 mL centrifugal filters (Merck Millipore) and stored at -80 °C.

### 2. Expression and purification of GRK1/2

GRK1 1–535(*38*) and GRK2 S670A(*47*) expression was performed in insect cells as described in Sterne-Marr *et al.*(*48*) The used plasmids were generously provided by the Tesmer Lab and the two constructs were chosen for the following reason: GRK1 was C-terminally truncated to prevent farnesylation of the kinase and concomitant membrane recruitment. In this way, neither GRK1 nor GRK2 were recruited to the membrane, therefore allowing for a uniform assay for both kinases. Note that GRK2 is recruited to the membrane via binding to the Gβγ subunit, which was absent in our simplified assay. The point-mutation in GRK2 was introduced as phosphorylation of the site was reported to shift specificity of the kinase towards non-receptor substrates(*47*), which is undesirable in our assay. All purification steps were performed either on ice or at 4 °C and all fractions were monitored by HPLC analysis using an 1100 Series system (Agilent), equipped with a POROS R1/10 4.6x100 C8 column (Applied Biosystems), running a gradient between buffer HPLC-A (ddH_2_O, 0.1% TFA) and HPLC-B (90% ACN, 10% ddH_2_O, 0.1% TFA). The cell pellet was resuspended in 5 mL washing buffer (50 mM TRIS pH 8, 300 mM NaCl, 5 mM imidazole, 10% v/v glycerol, 1 mM DTT) per 1 g pellet and DNaseI as well as 1 mM PMSF were added. The suspension was first treated 3 times for 30 s with an IKA T18 disperser (TURRAX), equipped with a S18N-19G dispersing tool, set to 10,000 rpm followed by 3 times 30 s of sonication (35% power, 50% cycles) using a Sonoplus HD 2070 connected to a UW 2070 (Bandelin electronic). Between every lysis step the suspension was cooled for 1 min on ice. Cell debris was separated by centrifugation at 35,000 g for 2–3 h and vacuum filtration of the cleared lysate through 0.45 µm nitrocellulose membranes (Merck, MF-Millipore). Ni-affinity chromatography on a 20 mL HisPrep FF 16/10 column (GE) with 140 mL washing buffer and 60 mL elution buffer (50 mM TRIS pH 8, 300 mM NaCl, 250 mM imidazole, 10% v/v glycerol, 1 mM DTT) was performed as a first purification step. The resulting protein solution was concentrated in 50 kDa Amicon-Ultra 15 mL centrifugal filters (Merck Millipore) and subsequently injected on a Superdex 200 Increase 10/600 column (GE) to obtain a monodisperse fraction in 70 mM TRIS pH 7.2, 150 mM NaCl, 1 mM DTT. Finally, the purified protein was concentrated in 50 kDa Amicon-Ultra 15 mL centrifugal filters (Merck Millipore) and stored at -80 °C.

### 3. Backbone Assignment Experiments

Backbone atoms of phosphorylated GPCR C-termini were assigned to their corresponding NMR signals by standard ^1^H,^13^C,^15^N triple resonance experiments performed on the GPCRs’ phosphorylated C-terminal peptides. Phosphorylated ^13^C,^15^N labeled GPCR C-termini were obtained by phosphorylation with GRK1 or GRK2. 300–600 µL samples containing 20–25 µM GRK1, 10 mM ATP and 300– 400 µM ^13^C,^15^N labeled GPCR C-terminus in 70 mM TRIS pH 7.2, 25 mM MgCl_2_, 2 mM TCEP were incubated for 40–48 h at 37 °C and 180 rpm in a TH 15 shaker (Edmund Bühler GmbH). The peptide concentration was chosen sufficiently high to obtain a phosphopeptide concentration of at least 500 µM in the final 160 µL NMR sample. Successful phosphorylation was confirmed prior to sample processing by recording a [^15^N,^1^H]-HMQC spectrum of the reaction mixture. To separate the phosphopeptide solution from the formed precipitate, the samples were centrifuged at 16,100 rcf and 4 °C for 10 min. Subsequently, the buffer of the supernatant was exchanged to 20 mM citrate pH 5.5, 150 mM NaCl, 1 mM DTT on a 1 mL PD MidiTrap G-25 (GE). The sample was concentrated to 140 µL in 2 kDa vivaspin 2 (Sartorius Stedim Biotech, rhodopsin C-terminus) or 3 kDa 0.5 mL Amicon-Ultra (Merck Millipore, βAR C-termini) centrifugal filters and 16–20 µL D_2_O were added to obtain a final volume of at least 160 µl.

The following spectra of the phosphopeptide were recorded in 3 mm tubes (Norell) at 298.1 K on a 700 MHz Bruker spectrometer, equipped with a CP-TCI 1H-13C/15N-2H ZGrad (βAR C-termini) or CP-TCI 1H&19F-13C/15N-2H XYZGrad (rhodopsin C-terminus) probe head. 1D ^1^H (ns = 128), XL-ALSOFAST-[^13^C,^1^H]-HMQC(*49*) (ns = 16 (βAR) or 32 (Rho), td(13C) = 280 (βAR) or 704 (Rho), SW(1H) = 16.2 ppm, o1p = 4.7 ppm, SW(13C) = 80 ppm, o2p = 42 ppm, d1 = 0.5 s, d21 = 1.7 ms, d22 = 1.7 ms), [^15^N,^1^H]-HSQC (ns = 4, SW(1H) = 16.2 ppm, o1p = 4.7 ppm, SW(15N) = 36 ppm, o2p = 118 ppm), HNCACB (from Bruker library, ns = 8 (βAR) or 16 (Rho), td(13C) = 176, td(15N) = 68) as well as CBCAcoNH (from Bruker library, ns = 8, td(13C) = 98, td(15N) = 68). The spectral width and offset of the triple resonance experiments was adjusted depending on the C-terminus as listed in supplementary table 1. Data were processed in Topspin 4.0.7. Backbone assignment of the phosphopeptides was performed in CARA (1.9.1.7). It is important to mention that phosphorylation of close-by sites can lead to a second state of the initial site. This influence is not restricted to one neighbor, possibly leading to several peaks. Nevertheless, assignment of these peaks could be done based on the triple resonance experiments using CARA (1.9.17).

The chemical shifts of ^1^H,^15^N-correlation signals of phosphorylated residues are highly pH sensitive. A pH-titration was performed to enable the transfer of assignments to spectra measured at other pH-values (Supplementary Figure 1). The experiments were either performed with the sample already used for the triple resonance experiments or a new, ^15^N labeled sample was prepared. A series of 1D 1H (ns = 128) and [^15^N,^1^H]-HMQC spectra (ns = 8, td(15N) = 256 (β_1_AR) or 512 (Rho, β_2_AR), SW(1H) = 16 ppm, o1p = 4.7 ppm, SW(15N) = 36 ppm, o2p = 118 ppm) was recorded at pH-values between 5.5 and 7.2 on 700 MHz (CP-TCI 1H-13C/15N-2H ZGrad probe head), 750 MHz (TXI 1H-13C/15N-2H ZGrad probe head) or 900 MHz (CP-TCI 1H-13C/15N-2H ZGard probe head) Bruker spectrometers.

### 4. Phosphorylation Assay

100 µM of the respective ^15^N-labeled C-terminus was mixed with 5 µM GRK1 or GRK2. As a last step prior to the measurement, 10 mM ATP was added to the solution. Reactions were performed in 70 mM TRIS pH 7.2, 25 mM MgCl_2_ and 2 mM TCEP in a total volume of 160 µL. Additionally, an inactive reference sample was pipetted, where the respective kinase was replaced by buffer.

ALSOFAST-[^15^N,^1^H]-HMQC experiments (ns = 16, td(15N) = 256, SW(1H) = 15.6 ppm, o1p = 4.7 ppm, SW(15N) = 36 ppm, o2p = 120, d1 = 0.2 s) of the samples were measured on a 500 MHz Bruker spectrometer (CP-QCI 1H & 19F/31P-13C/15N/2H Z-Grad probe head) in 3 mm tubes (Norell) at 298.1 K every 2–6 h over a period of 30–60 h. At the end of the measurement of the time-series the pH for both samples (reference and time-series sample) was dropped to pH 5.5 for normalization purposes.

After data processing in Topspin 4.0.7, signal integrals were read out in Julia 1.9.2 using a previously described script(*50*). Data was normalized according to the procedure described by Theillet *et al*.(*35*) and further processed in JupyterLab 4.0.7. A list of used reporter peaks as well as a ranking of the individual 100% phosphorylation levels can be found in supplementary table 6. Fits of biphasic curves were generated according to the equations 1.1–1.5 (Supplementary Figure 1) in JupyterLab 4.0.7. For data shown in figure 4, signals were read out, normalized by peaks which are unaffected by the phosphorylation reaction and further processed in Julia 1.9.2 using a previously described script(*50*).

### 5. Split luciferase assay

The split luciferase assay was performed based on Littmann *et al.*(*33*) and Jaiswal.(*51*) Briefly, 1.6 × 10^6^ quadruple GRK knockout cells (ΔQ-GRK, ΔGRK2/3/5/6) were seeded in 21 cm^2^ dishes and transfected the following day with 4 µg Gα_q_-PLCβ3 split luciferase sensor containing Gα_q_ subunit fused to the small part of the click-beetle luciferase and phospholipase Cβ3 (PLCβ3) fused to the large part the click-beetle luciferase, 2 µg of muscarinic acetylcholine receptor M1 (M1mAChR) and 3 µg of an individual GRK or empty vector. All transfections were conducted following the Effectene transfection reagent manual by Qiagen (#301427) and then incubated at 37 °C overnight. 60,000 cells were seeded per well into poly-D-lysine-coated 96-well plates (Brand, 781965). After 24 h, the cells were washed twice with measuring buffer (140 mM NaCl, 10 mM HEPES, 5.4 mM KCl, 2 mM CaCl_2_, 1 mM MgCl_2_; pH 7.3) and 1 nM D-luciferin (88293, Thermo Fisher Scientific) was added in measuring buffer. A Synergy Neo2 plate reader (Biotek), operated with the Gen5 software (version 2.09), with a custom-made filter (excitation bandwidth 541–550 nm, emission 560–595 nm, fluorescence filter 620/15 nm) was used to perform the measurements. The baseline was monitored for 7.5 min. After the addition of acetylcholine (ACh; Sigma-Aldrich I5627, in water), the measurements were continued for 27 minutes. Luminescence values after ligand stimulation were corrected for non-specific signal by using values from vehicle stimulation and analyzed over time to determine the time-points showing stable and maximum luminescence. An average of responses from the last three time-points were used for concentration-response curves. Calculations were conducted using Excel 2016. Further data processing was done in GraphPad Prism 7.03. with *n* = 4 independent experiments.

### 6. Intermolecular bioluminescence resonance energy transfer (BRET)

The GRK-selective β-arrestin recruitment assay was performed as described in Drube *et al.*(*34*) 1.6 × 10^6^ ΔQ-GRK cells were seeded in 21 cm^2^ dishes and transfected the next day with 0.5 μg of human β_2_ adrenergic receptor (β_2_AR) C-terminally fused to Nano luciferase (NanoLuc), 1 μg of β-arrestin constructs N-terminally fused to a Halo-ligand binding Halo-Tag and 0.25 μg of one GRK or empty vector. All transfections were conducted following the Effectene transfection reagent manual by Qiagen (#301427) and then incubated at 37 °C overnight. Into poly-D-lysine-coated 96-well plates (Brand, 781965), 40,000 cells were seeded per well in presence of Halo-ligand (Promega, G980A) at a ratio of 1:2,000. Each transfection was seeded in three technical replicates and one mock labelling condition without the addition of the Halo-ligand. After 24 h, the cells were washed twice with measuring buffer (140 mM NaCl, 10 mM HEPES, 5.4 mM KCl, 2 mM CaCl_2_, 1 mM MgCl_2_; pH 7.3) and NanoLuc-substrate furimazine (Promega, N157B) was added in a ratio of 1:35,000 in measuring buffer. A Synergy Neo2 plate reader (Biotek), operated with the Gen5 software (version 2.09), with a custom-made filter (excitation bandwidth 541–550 nm, emission 560–595 nm, fluorescence filter 620/15 nm) was used to perform the measurements. The baseline was monitored for 3 min. After the addition of different isoproterenol concentrations as indicated (Iso; Sigma-Aldrich I5627, in water), the measurements were continued for 15 minutes. To obtain the BRET values, the acceptor (Halo-tag) signal was divided by the donor (NanoLuc) signal and the mean of the technical replicates was calculated. By subtracting the values measured for the respective mock labelling conditions, the initial BRET values were corrected for labelling efficiency. For concentration-response curves, Halo-corrected BRET changes were calculated by the division of the corrected and averaged values measured after ligand stimulation (timepoints 2–5 min after stimulation) by the respective corrected and averaged baseline values. Subsequently, this corrected BRET change was divided by the vehicle (buffer-stimulated) control for the final dynamic Δ net BRET change. These calculations were conducted using Excel 2016. Curves and corresponding SEM for concentration-dependent β-arrestin recruitment were calculated with GraphPad Prism 7.03. using data from *n* = 4 independent experiments.

## Acknowledgements

The authors would like to thank Carla Ferreira Rodrigues and Lucas Kirchen for help with assignments of the peptides. Further, the authors would like to acknowledge Christopher Waudby for excellent help with the program Julia. The authors would also like to thank the Tesmer Lab for generously providing the GRK plasmids. A.L., N.L. and P.R. thank the Biomolecular Structure and Mechanism PhD Program of the Life Science Zurich Graduate School. The authors are grateful to Frédéric H.T. Allain for sharing lab space and equipment. This work was supported by the Swiss National Science Foundation by grant 31-208029 and by ETH Zürich with project grant ETH-37 19-2 to A.D.G.

## Supplementary material

Details on resonance assignment, analysis of phosphorylation rates, and phosphorylation patterns in β_2_AR Cterm peptides of different length are available as supplementary material. Supplementary Tables 1 – 6

Supplementary Figures 1 – 5

## References

1. A. S. Hauser, M. M. Attwood, M. Rask-Andersen, H. B. Schiöth, D. E. Gloriam, Trends in GPCR drug discovery: new agents, targets and indications. Nat. Rev. Drug Discov. 16, 829–842 (2017).

2. R. T. Premont, R. R. Gainetdinov, Physiological Roles of G Protein–Coupled Receptor Kinases and Arrestins. Annu. Rev. Physiol. 69, 511–534 (2007).

3. A. B. Tobin, A. J. Butcher, K. C. Kong, Location, location, location…site-specific GPCR phosphorylation offers a mechanism for cell-type-specific signalling. Trends Pharmacol. Sci. 29, 413–420 (2008).

4. K. N. Nobles, K. Xiao, S. Ahn, A. K. Shukla, C. M. Lam, S. Rajagopal, R. T. Strachan, T.-Y. Huang, E. A. Bressler, M. R. Hara, S. K. Shenoy, S. P. Gygi, R. J. Lefkowitz, Distinct phosphorylation sites on the β(2)-adrenergic receptor establish a barcode that encodes differential functions of β-arrestin. Sci. Signal. 4, ra51 (2011).

5. R. H. Oakley, S. A. Laporte, J. A. Holt, L. S. Barak, M. G. Caron, Association of β-Arrestin with G Protein-coupled Receptors during Clathrin-mediated Endocytosis Dictates the Profile of Receptor Resensitization. J. Biol. Chem. 274, 32248–32257 (1999).

6. X. E. Zhou, Y. He, P. W. de Waal, X. Gao, Y. Kang, N. Van Eps, Y. Yin, K. Pal, D. Goswami, T. A. White, A. Barty, N. R. Latorraca, H. N. Chapman, W. L. Hubbell, R. O. Dror, R. C. Stevens, V. Cherezov, V. V. Gurevich, P. R. Griffin, O. P. Ernst, K. Melcher, H. E. Xu, Identification of Phosphorylation Codes for Arrestin Recruitment by G Protein-Coupled Receptors. Cell 170, 457–469.e13 (2017).

7. J. Maharana, P. Sarma, M. K. Yadav, S. Saha, V. Singh, S. Saha, M. Chami, R. Banerjee, A. K. Shukla, Structural snapshots uncover a key phosphorylation motif in GPCRs driving β-arrestin activation. Mol. Cell 83, 2091–2107.e7 (2023).

8. P. Isaikina, I. Petrovic, R. P. Jakob, P. Sarma, A. Ranjan, M. Baruah, V. Panwalkar, T. Maier, A. K. Shukla, S. Grzesiek, A key GPCR phosphorylation motif discovered in arrestin2⋅CCR5 phosphopeptide complexes. Mol. Cell 83, 2108–2121.e7 (2023).

9. A. Sente, R. Peer, A. Srivastava, M. Baidya, A. M. Lesk, S. Balaji, A. K. Shukla, M. M. Babu, T. Flock, Molecular mechanism of modulating arrestin conformation by GPCR phosphorylation. Nat. Struct. Mol. Biol. 25, 538–545 (2018).

10. Q. Chen, J. J. G. Tesmer, G protein–coupled receptor interactions with arrestins and GPCR kinases: The unresolved issue of signal bias. J. Biol. Chem. 298, 102279 (2022).

11. J. A. Pitcher, Z. L. Fredericks, W. C. Stone, R. T. Premont, R. H. Stoffel, W. J. Koch, R. J. Lefkowitz, Phosphatidylinositol 4,5-Bisphosphate (PIP2)-enhanced G Protein-coupled Receptor Kinase (GRK) Activity: Location, Structure, and regulation of the PIP2 binding site distinguishes the GRK subfamilies. J. Biol. Chem. 271, 24907–24913 (1996).

12. J. A. Pitcher, N. J. Freedman, R. J. Lefkowitz, G protein-coupled receptor kinases. Annu. Rev. Biochem. 67, 653–692 (1998).

13. C. V. Carman, L. S. Barak, C. Chen, L.-Y. Liu-Chen, J. J. Onorato, S. P. Kennedy, M. G. Caron, J. L. Benovic, Mutational Analysis of Gβγ and Phospholipid Interaction with G Protein-coupled Receptor Kinase 2 . J. Biol. Chem. 275, 10443–10452 (2000).

14. L. R. Pearce, D. Komander, D. R. Alessi, The nuts and bolts of AGC protein kinases. Nat. Rev. Mol. Cell Biol. 11, 9–22 (2010).

15. E. S. F. Matthees, J. C. Filor, N. Jaiswal, M. Reichel, N. Youssef, G. D’Uonnolo, M. Szpakowska, J. Drube, G. M. König, E. Kostenis, A. Chevigné, A. Godbole, C. Hoffmann, GRK specificity and Gβγ dependency determines the potential of a GPCR for arrestin-biased agonism. *Commun*. Biol. 7, 1–12 (2024).

16. J. Li, A. Inoue, A. Manglik, M. von Zastrow, Role of the G protein-coupled receptor kinase 2/3 N terminus in discriminating the endocytic effects of opioid agonist drugs. Mol. Pharmacol. 107, 100003 (2025).

17. I. Torrecilla, E. J. Spragg, B. Poulin, P. J. McWilliams, S. C. Mistry, A. Blaukat, A. B. Tobin, Phosphorylation and regulation of a G protein–coupled receptor by protein kinase CK2. J. Cell Biol. 177, 127–137 (2007).

18. J. M. Busillo, S. Armando, R. Sengupta, O. Meucci, M. Bouvier, J. L. Benovic, Site-specific Phosphorylation of CXCR4 Is Dynamically Regulated by Multiple Kinases and Results in Differential Modulation of CXCR4 Signaling. J. Biol. Chem. 285, 7805–7817 (2010).

19. A. J. Butcher, R. Prihandoko, K. C. Kong, P. McWilliams, J. M. Edwards, A. Bottrill, S. Mistry, A. B. Tobin, Differential G-protein-coupled Receptor Phosphorylation Provides Evidence for a Signaling Bar Code. J. Biol. Chem. 286, 11506–11518 (2011).

20. M. Trester-Zedlitz, A. Burlingame, B. Kobilka, M. von Zastrow, Mass Spectrometric Analysis of Agonist Effects on Posttranslational Modifications of the β-2 Adrenoceptor in Mammalian Cells. Biochemistry 44, 6133–6143 (2005).

21. S. Wu, M. Birnbaumer, Z. Guan, Phosphorylation Analysis of G Protein-Coupled Receptor by Mass Spectrometry: Identification of a Phosphorylation Site in V2 Vasopressin Receptor. Anal. Chem. 80, 6034–6037 (2008).

22. A. J. Butcher, B. D. Hudson, B. Shimpukade, E. Alvarez-Curto, R. Prihandoko, T. Ulven, G. Milligan, A. B. Tobin, Concomitant Action of Structural Elements and Receptor Phosphorylation Determines Arrestin-3 Interaction with the Free Fatty Acid Receptor FFA4. J. Biol. Chem. 289, 18451–18465 (2014).

23. J. Kaufmann, N. K. Blum, F. Nagel, A. Schuler, J. Drube, C. Degenhart, J. Engel, J. E. Eickhoff, P. Dasgupta, S. Fritzwanker, M. Guastadisegni, C. Schulte, E. Miess-Tanneberg, H. M. Maric, M. Spetea, A. Kliewer, M. Baumann, B. Klebl, R. K. Reinscheid, C. Hoffmann, S. Schulz, A bead-based GPCR phosphorylation immunoassay for high-throughput ligand profiling and GRK inhibitor screening. *Commun*. Biol. 5, 1206 (2022).

24. H. Ohguro, K. Palczewski, L. H. Ericsson, K. A. Walsh, R. S. Johnson, Sequential phosphorylation of rhodopsin at multiple sites. Biochemistry 32, 5718–5724 (1993).

25. H. Ohguro, J. P. V. Hooser, A. H. Milam, K. Palczewski, Rhodopsin Phosphorylation and Dephosphorylation in Vivo. J. Biol. Chem. 270, 14259–14262 (1995).

26. L. Zhang, C. D. Sports, S. Osawa, E. R. Weiss, Rhodopsin Phosphorylation Sites and Their Role in Arrestin Binding . J. Biol. Chem. 272, 14762–14768 (1997).

27. M. J. Kennedy, K. A. Lee, G. A. Niemi, K. B. Craven, G. G. Garwin, J. C. Saari, J. B. Hurley, Multiple Phosphorylation of Rhodopsin and the In Vivo Chemistry Underlying Rod Photoreceptor Dark Adaptation. Neuron 31, 87–101 (2001).

28. A. W. Azevedo, T. Doan, H. Moaven, I. Sokal, F. Baameur, S. A. Vishnivetskiy, K. T. Homan, J. J. Tesmer, V. V. Gurevich, J. Chen, F. Rieke, C-terminal threonines and serines play distinct roles in the desensitization of rhodopsin, a G protein-coupled receptor. eLife 4, e05981 (2015).

29. S. Rajagopal, S. K. Shenoy, GPCR desensitization: Acute and prolonged phases. Cell. Signal. 41, 9–16 (2018).

30. D. T. Lodowski, J. A. Pitcher, W. D. Capel, R. J. Lefkowitz, J. J. G. Tesmer, Keeping G Proteins at Bay: A Complex Between G Protein-Coupled Receptor Kinase 2 and Gßγ. Science 300, 1256– 1262 (2003).

31. E. V. Gurevich, J. J. G. Tesmer, A. Mushegian, V. V. Gurevich, G protein-coupled receptor kinases: more than just kinases and not only for GPCRs. Pharmacol. Ther. 133, 40–69 (2012).

32. V. M. Tesmer, T. Kawano, A. Shankaranarayanan, T. Kozasa, J. J. G. Tesmer, Snapshot of Activated G Proteins at the Membrane: The Gαq-GRK2-Gβγ Complex. Science 310, 1686–1690 (2005).

33. T. Littmann, T. Ozawa, C. Hoffmann, A. Buschauer, G. Bernhardt, A split luciferase-based probe for quantitative proximal determination of Gαq signalling in live cells. Sci. Rep. 8, 17179 (2018).

34. J. Drube, R. S. Haider, E. S. F. Matthees, M. Reichel, J. Zeiner, S. Fritzwanker, C. Ziegler, S. Barz, L. Klement, J. Filor, V. Weitzel, A. Kliewer, E. Miess-Tanneberg, E. Kostenis, S. Schulz, C. Hoffmann, GPCR kinase knockout cells reveal the impact of individual GRKs on arrestin binding and GPCR regulation. Nat. Commun. 13, 540 (2022).

35. F.-X. Theillet, H. M. Rose, S. Liokatis, A. Binolfi, R. Thongwichian, M. Stuiver, P. Selenko, Site-specific NMR mapping and time-resolved monitoring of serine and threonine phosphorylation in reconstituted kinase reactions and mammalian cell extracts. Nat. Protoc. 8, 1416–1432 (2013).

36. E. S. F. Matthees, R. S. Haider, C. Hoffmann, J. Drube, Differential Regulation of GPCRs—Are GRK Expression Levels the Key? Front. Cell Dev. Biol. 9 (2021).

37. C. Huang, K. Yoshino-Koh, J. J. G. Tesmer, A Surface of the Kinase Domain Critical for the Allosteric Activation of G Protein-coupled Receptor Kinases. J. Biol. Chem. 284, 17206–17215 (2009).

38. P. Singh, B. Wang, T. Maeda, K. Palczewski, J. J. G. Tesmer, Structures of Rhodopsin Kinase in Different Ligand States Reveal Key Elements Involved in G Protein-coupled Receptor Kinase Activation. J. Biol. Chem. 283, 14053–14062 (2008).

39. Q. Chen, M. Plasencia, Z. Li, S. Mukherjee, D. Patra, C.-L. Chen, T. Klose, X.-Q. Yao, A. A. Kossiakoff, L. Chang, P. C. Andrews, J. J. G. Tesmer, Structures of rhodopsin in complex with G-protein-coupled receptor kinase 1. Nature 595, 600–605 (2021).

40. J. J. Onorato, K. Palczewski, J. W. Regan, M. G. Caron, R. J. Lefkowitz, J. L. Benovic, Role of acidic amino acids in peptide substrates of the beta-adrenergic receptor kinase and rhodopsin kinase. Biochemistry 30, 5118–5125 (1991).

41. J. L. Johnson, T. M. Yaron, E. M. Huntsman, A. Kerelsky, J. Song, A. Regev, T.-Y. Lin, K. Liberatore, D. M. Cizin, B. M. Cohen, N. Vasan, Y. Ma, K. Krismer, J. T. Robles, B. van de Kooij, A. E. van Vlimmeren, N. Andrée-Busch, N. F. Käufer, M. V. Dorovkov, A. G. Ryazanov, Y. Takagi, E. R. Kastenhuber, M. D. Goncalves, B. D. Hopkins, O. Elemento, D. J. Taatjes, A. Maucuer, A. Yamashita, A. Degterev, M. Uduman, J. Lu, S. D. Landry, B. Zhang, I. Cossentino, R. Linding, J. Blenis, P. V. Hornbeck, B. E. Turk, M. B. Yaffe, L. C. Cantley, An atlas of substrate specificities for the human serine/threonine kinome. Nature 613, 759–766 (2023).

42. L. Hinz, A. Ahles, B. Ruprecht, B. Küster, S. Engelhardt, Two serines in the distal C-terminus of the human ß1-adrenoceptor determine ß-arrestin2 recruitment. PLOS ONE 12, e0176450 (2017).

43. Y. Shiraishi, M. Natsume, Y. Kofuku, S. Imai, K. Nakata, T. Mizukoshi, T. Ueda, H. Iwaï, I. Shimada, Phosphorylation-induced conformation of β2-adrenoceptor related to arrestin recruitment revealed by NMR. Nat. Commun. 9, 194 (2018).

44. D. Mayer, F. F. Damberger, M. Samarasimhareddy, M. Feldmueller, Z. Vuckovic, T. Flock, B. Bauer, E. Mutt, F. Zosel, F. H. T. Allain, J. Standfuss, G. F. X. Schertler, X. Deupi, M. E. Sommer, M. Hurevich, A. Friedler, D. B. Veprintsev, Distinct G protein-coupled receptor phosphorylation motifs modulate arrestin affinity and activation and global conformation. Nat. Commun. 10, 1261 (2019).

45. R. Tatsumi, S. Aihara, S. Matsune, J. Aoki, A. Inoue, T. Shimizu, M. Nakamura, Stepwise phosphorylation of BLT1 defines complex assemblies with β-arrestin serving distinct functions. FASEB J. Off. Publ. Fed. Am. Soc. Exp. Biol. 37, e23213 (2023).

46. R. S. Haider, E. S. F. Matthees, J. Drube, M. Reichel, U. Zabel, A. Inoue, A. Chevigné, C. Krasel, X. Deupi, C. Hoffmann, β-arrestin1 and 2 exhibit distinct phosphorylation-dependent conformations when coupling to the same GPCR in living cells. Nat. Commun. 13, 5638 (2022).

47. J. A. Pitcher, J. J. G. Tesmer, J. L. R. Freeman, W. D. Capel, W. C. Stone, R. J. Lefkowitz, Feedback Inhibition of G Protein-coupled Receptor Kinase 2 (GRK2) Activity by Extracellular Signal-regulated Kinases. J. Biol. Chem. 274, 34531–34534 (1999).

48. R. Sterne-Marr, A. I. Baillargeon, K. R. Michalski, J. J. G. Tesmer, Expression, purification, and analysis of G-protein-coupled receptor kinases. Methods Enzymol. 521, 347–366 (2013).

49. P. Rößler, D. Mathieu, A. D. Gossert, Enabling NMR Studies of High Molecular Weight Systems Without the Need for Deuteration: The XL-ALSOFAST Experiment with Delayed Decoupling. Angew. Chem. Int. Ed. 59, 19329–19337 (2020).

50. C. A. Waudby, S. Alvarez-Teijeiro, E. Josue Ruiz, S. Suppinger, N. Pinotsis, P. R. Brown, A. Behrens, J. Christodoulou, A. Mylona, An intrinsic temporal order of c-JUN N-terminal phosphorylation regulates its activity by orchestrating co-factor recruitment. Nat. Commun. 13, 6133 (2022).

51. N. Jaiswal, New insights into non-canonical desensitization of Gq signaling by GRK2/3 expression levels (2023). https://nbn-resolving.org/urn:nbn:de:gbv:27-dbt-20231115-151703-000.

