## Supplementary material for "GRKs phosphorylate GPCR C-terminal peptides in a hierarchical manner"

This .pdf includes:

**Supplementary Table 1.** Spectral width and offset used for HNCACB and CBCAcoNH experiments on the phosphorylated C-termini. Parameters are defined as in the experiments from the Bruker library. All values are given in ppm.

| C-terminus | SW($^1$H) | o1p | SW($^{15}$N) | o2p | SW($^{13}$C) | o3p |
|---|---|---|---|---|---|---|
| **Rho** | 15.5 | 4.7 | 20 | 121.5 | 61 | 46.5 |
| **$\beta_1$AR** | 16.2 | 4.69 | 21 | 118 | 52 | 48 |
| **$\beta_2$AR** | 16.2 | 4.69 | 21 | 118 | 52 | 45 |

**Supplementary Table 2.** Assigned amide-H, amide-N, Cα and Cβ chemical shifts (in ppm) of Rho Cterm.

| Residue | $\delta_H$ | $\delta_N$ | $\delta_{C\alpha}$ | $\delta_{C\beta}$ |
|---|---|---|---|---|
| 330 | 8.6268 | 120.6364 | 54.4890 | 40.9040 |
| 331 | 8.4323 | 120.1101 | 54.4760 | 40.8903 |
| 332 | 8.2692 | 120.9537 | 56.6317 | 29.9075 |
| 333 | 8.2187 | 124.4825 | 52.5663 | 18.9804 |
| 334 | 8.1871 | 114.8700 | 58.3016 | 63.5313 |
| 335 | 8.1076 | 115.6793 | 61.8521 | 69.5041 |
| 336 | 8.0758 | 117.2631 | 62.0963 | 69.4906 |
| 337 | 8.1071 | 123.0976 | 62.1080 | 32.5572 |
| 338 | 8.3567 | 120.0396 | 58.0567 | 63.5511 |
| 339 | 8.3692 | 123.9799 | 56.1509 | 33.0375 |
| 340 | 8.1863 | 115.6430 | 61.8742 | 69.7167 |
| 341 | 8.4084 | 123.4529 | 56.3791 | 30.1721 |
| 342 | 8.1742 | 115.0464 | 61.6298 | 69.7335 |
| 343 | 8.6816 | 118.1808 | 57.5774 | 66.1637 |
| 344 | 8.2302 | 121.7998 | 55.4344 | 29.4483 |
| 345 | 8.0600 | 121.5228 | 61.8679 | 32.5625 |
| 346 | 8.3506 | 129.6129 | 50.1936 | 18.0283 |
| 348 | 7.8813 | 129.9233 | 53.5267 | 19.9321 |

**Supplementary Table 3.** Assigned amide-H, amide-N, Cα and Cβ chemical shifts (in ppm) of $\beta_1$AR Cterm.

| Residue | $\delta_H$ | $\delta_N$ | $\delta_{C\alpha}$ | $\delta_{C\beta}$ |
|---|---|---|---|---|
| 407 | 8.6748 | 120.7247 | 54.4817 | |
| 408 | 8.0249 | 121.2335 | 54.8218 | 29.9601 |
| 410 | 8.4934 | 121.9041 | 56.2045 | |
| 411 | 8.4459 | 125.7780 | 52.6998 | 19.4510 |
| 412 | 8.2014 | 115.3094 | 58.6757 | 63.9192 |
| 413 | 8.4720 | 110.9107 | 45.5212 | |
| 414 | 8.2433 | 118.9514 | 58.6698 | 28.1878 |
| 415 | 8.3527 | 124.7956 | 55.4003 | 42.4493 |
| 416 | 8.2100 | 124.8725 | 52.4736 | 19.3948 |
| 417 | 8.2432 | 121.4873 | 53.9528 | 30.4818 |
| 425 | 8.4752 | 109.7322 | 45.3102 | 0.0000 |
| 426 | 8.0714 | 123.9221 | 52.4727 | 19.6951 |
| 427 | 8.1468 | 121.1868 | 52.6573 | 19.7094 |
| 428 | 8.2528 | 116.9659 | 60.1643 | 69.9442 |
| 430 | 8.3282 | 120.5576 | 54.6643 | 41.0837 |
| 431 | 8.2869 | 120.7946 | 54.4974 | 0.0000 |
| 432 | 8.2479 | 120.1966 | 54.5476 | 0.0000 |
| 433 | 8.2653 | 120.6594 | 54.5560 | 41.1427 |
| 434 | 8.3185 | 120.8032 | 54.6929 | 41.0513 |
| 435 | 8.0198 | 120.4072 | 62.7524 | 32.7409 |
| 436 | 8.2220 | 124.0666 | 63.1270 | |
| 437 | 8.4567 | 112.6485 | 45.3814 | |
| 438 | 8.1063 | 123.6890 | 52.4797 | 19.6083 |
| 439 | 8.2252 | 116.6906 | 60.0868 | 69.9652 |
| 443 | 8.2616 | 119.8860 | 56.1598 | |
| 444 | 8.2045 | 123.4091 | 55.2928 | 42.3998 |
| 446 | 8.2001 | 122.2930 | 54.4488 | 29.4890 |
| 448 | 7.8834 | 119.4718 | 57.1996 | 29.3540 |
| 449 | 8.0037 | 126.6220 | 52.9622 | 19.3827 |
| 450 | 7.5100 | 106.5052 | 45.4281 | 0.0000 |
| 451 | 8.2318 | 118.1133 | 55.7916 | 41.2546 |
| 452 | 8.6137 | 120.5088 | 53.5695 | 39.0371 |
| 453 | 8.4286 | 109.6718 | 45.7507 | 0.0000 |
| 454 | 8.2285 | 108.8290 | 45.3416 | 0.0000 |
| 455 | 8.1578 | 123.8561 | 52.6493 | 19.6240 |
| 456 | 8.3693 | 123.8363 | 52.7515 | 19.3825 |
| 457 | 8.2841 | 123.2265 | 52.7791 | 19.3428 |
| 458 | 8.2847 | 119.3163 | 54.5806 | 41.1250 |
| 459 | 8.1097 | 115.3413 | 58.9422 | 64.1290 |
| 460 | 8.4257 | 122.4071 | 54.6663 | 41.2129 |
| 461 | 8.2970 | 116.8271 | 58.8700 | 63.8979 |
| 462 | 8.4124 | 118.0603 | 59.0184 | 63.7721 |
| 463 | 8.1152 | 123.2063 | 55.4175 | 42.4488 |
| 464 | 8.1942 | 120.7627 | 54.4489 | 41.2568 |
| 465 | 8.1289 | 122.0299 | 54.5484 | 30.0358 |
| 466 | 0.0000 | 0.0000 | 63.4197 | 32.2037 |
| 467 | 8.4329 | 119.1904 | 58.5755 | |

| | | | | |
|---|---|---|---|---|
| 468 | 8.3967 | 124.8731 | 54.2578 | 30.3631 |
| 469 | 0.0000 | 0.0000 | 63.5075 | 32.1532 |
| 470 | 8.5281 | 109.7191 | 45.3332 | 0.0000 |
| 471 | 8.0406 | 120.0912 | 58.0961 | 39.9286 |
| 472 | 8.2763 | 125.7296 | 52.6154 | 19.4497 |
| 473 | 8.3416 | 115.3001 | 58.3438 | 64.2313 |
| 474 | 8.4476 | 122.6984 | 56.7867 | 30.3540 |
| 475 | 8.3264 | 117.0001 | 58.5305 | 63.9418 |
| 476 | 8.3959 | 124.4956 | 56.4792 | 33.1021 |
| 477 | 7.7455 | 125.1442 | 63.7412 | 33.3745 |

**Supplementary Table 4.** Assigned amide-H, amide-N, Cα and Cβ chemical shifts (in ppm) of $\beta_2$AR Cterm.

| Residue | $\delta_H$ | $\delta_N$ | $\delta_{C\alpha}$ | $\delta_{C\beta}$ |
|---|---|---|---|---|
| 342 | 8.5051 | 121.6949 | 55.2513 | 42.5579 |
| 343 | 8.4526 | 122.3174 | 55.8609 | 30.6550 |
| 344 | 8.4534 | 123.5878 | 56.2277 | 30.7015 |
| 345 | 8.4117 | 117.5324 | 58.5990 | 63.6645 |
| 346 | 8.7293 | 118.5103 | 57.5801 | 66.0573 |
| 347 | 7.9889 | 123.3963 | 55.5038 | 41.9454 |
| 348 | 8.0978 | 122.1300 | 55.9487 | 32.7473 |
| 349 | 8.0459 | 124.4559 | 52.1429 | 19.0776 |
| 350 | 7.9397 | 118.7947 | 57.6691 | 38.7660 |
| 351 | 8.2132 | 110.0440 | 45.2382 | |
| 352 | 7.8237 | 123.7512 | 54.4609 | 40.6516 |
| 354 | 8.0395 | 119.6269 | 57.8571 | 38.8975 |
| 355 | 8.1274 | 117.5484 | 57.8726 | 63.8969 |
| 356 | 8.2801 | 117.8561 | 58.3961 | 63.6493 |
| 357 | 8.3696 | 120.2569 | 53.2075 | 38.7502 |
| 358 | 8.2424 | 108.8109 | 45.4427 | |
| 360 | 8.1746 | 113.8524 | 62.1686 | 69.4836 |
| 361 | 8.3617 | 110.9406 | 45.3101 | |
| 362 | 8.1947 | 120.6549 | 56.6929 | 30.0712 |
| 363 | 8.4112 | 121.0202 | 55.6906 | 29.0669 |
| 364 | 8.2767 | 116.7430 | 58.5220 | 63.6043 |
| 365 | 8.3130 | 110.4674 | 45.1061 | |
| 366 | 7.9389 | 120.2023 | 57.9084 | 38.6012 |
| 367 | 8.2361 | 121.8709 | 55.0514 | 29.2577 |
| 368 | 8.0503 | 121.8719 | 62.3707 | 32.7188 |
| 369 | 8.5175 | 124.4688 | 56.5701 | 29.8777 |
| 372 | 8.3129 | 121.5257 | 55.8821 | 29.6510 |
| 373 | 8.3977 | 122.0855 | 56.8914 | 29.8715 |
| 374 | 8.3847 | 120.0567 | 53.0339 | 38.6124 |
| 375 | 8.2138 | 122.1150 | 56.4538 | 32.7454 |
| 376 | 8.1137 | 122.4402 | 55.1586 | 42.0470 |
| 377 | 8.0505 | 122.4392 | 55.0569 | 42.0606 |
| 378 | 8.1846 | 119.7886 | 58.3154 | 27.9489 |
| 379 | 8.3984 | 123.0745 | 56.4848 | 30.4686 |
| 380 | 8.2349 | 121.2477 | 54.0504 | 41.0315 |
| 381 | 8.0749 | 123.4907 | 52.8272 | 41.6606 |
| 383 | 8.4321 | 109.3189 | 45.1093 | |
| 384 | 7.9157 | 112.9060 | 61.9629 | 69.6883 |
| 385 | 8.5165 | 122.7175 | 56.6856 | 29.8869 |
| 386 | 8.1783 | 120.7814 | 54.2360 | 40.9187 |
| 387 | 8.0387 | 120.4795 | 57.9051 | 39.3327 |
| 388 | 7.9405 | 121.9189 | 62.7758 | 32.4339 |
| 389 | 8.0601 | 111.2636 | 45.1098 | |
| 390 | 8.2348 | 118.0809 | 55.4726 | 29.2696 |
| 391 | 8.4560 | 121.7532 | 55.8723 | 29.3995 |
| 392 | 8.4075 | 110.2766 | 45.1061 | |
| 393 | 8.0240 | 114.1740 | 61.7626 | 69.8895 |

| | | | | |
|---|---|---|---|---|
| 394 | 8.2069 | 124.0206 | 59.6847 | 32.4573 |
| 396 | 8.3034 | 115.7142 | 58.3100 | 63.7982 |
| 397 | 8.2473 | 121.9008 | 54.2476 | 41.0428 |
| 398 | 8.2463 | 118.5703 | 53.2354 | 38.8101 |
| 399 | 7.9931 | 120.8504 | 60.7483 | 38.8106 |
| 400 | 8.3560 | 123.8021 | 54.2163 | 40.9868 |
| 401 | 8.1906 | 116.1668 | 58.9249 | 63.3921 |
| 402 | 8.3836 | 121.0228 | 55.8786 | 28.8611 |
| 403 | 8.1331 | 108.7775 | 45.7356 | |
| 404 | 8.1794 | 120.5643 | 56.0817 | 30.4946 |
| 405 | 8.5000 | 119.7763 | 53.2351 | 38.6147 |
| 406 | 8.2916 | 119.6398 | 58.3203 | 28.0413 |
| 407 | 8.4365 | 118.2008 | 58.5138 | 63.7008 |
| 408 | 8.1254 | 115.3337 | 61.7677 | 69.5296 |
| 409 | 8.3446 | 120.6155 | 53.2338 | 38.8121 |
| 410 | 8.2375 | 121.0082 | 54.6437 | 40.9335 |
| 411 | 8.1185 | 115.4909 | 58.5108 | 63.5886 |
| 412 | 8.1403 | 124.0412 | 55.0625 | 41.9151 |
| 413 | 7.6075 | 127.7084 | 56.4803 | 43.2817 |

**Supplementary Table 5. Rate constants derived from fitted curves.** (**a**) Rate constants for individual phosphorylation sites of logarithmic curves of two individual datasets (1,2). Integrals were calculated using previously described scripts in Julia 1.9.2.[21] Curves were fitted using JupyterLab4.0.7. (**b**) Rate constants of individual phosphorylation sites of biphasic curves. Values were determined based on equations 1.1–1.5 using JupyterLab4.0.7. The integral at the starting point of the phosphorylation reaction was included into the fit ($UU_0$). Note: Rate constants are shown for completeness. As this assay is conducted based on the basal activity of the kinase, the described values are note representative of the reaction rates in cells.

**a**

| | | | | | | |
|---|---|---|---|---|---|---|
| **Rho [$10^{-3}$/h]** | | **pS334** | **pT336** | **pS338** | **pT340** | **pS343** |
| | 1 | 6 | 42 | - | - | 15 |
| | 2 | 13 | 21 | - | - | 20 |
| **$\beta_1$AR [$10^{-3}$/h]** | | **pS412** | **pS473** | **pS459** | **pS461** | **pS462** |
| | 1 | 18 | 30 | 25 | 39 | - |
| | 2 | 27 | 29 | - | 38 | - |
| **$\beta_2$AR [$10^{-3}$/h]** | | **pS345** | **pS346** | **pS364** | | |
| | 1 | 17 | 16 | 29 | | |
| | 2 | 24 | 27 | 38 | | |

**b**

| **Rho** | | | **T340** | |
|---|---|---|---|---|
| | | **$k_1$** | **$k_2$** | **$UU_0$** |
| **S343 [$10^{-3}$/h]** | 1 | 23 | 24 | 110 |
| | 2 | 30 | 10 | 83 |

| **Rho** | | | **S338** | |
|---|---|---|---|---|
| | | **$k_1$** | **$k_2$** | **$UU_0$** |
| **S343 [$10^{-3}$/h]** | 1 | 19 | 91 | 126 |
| | 2 | 35 | 57 | 78 |

| **$\beta_1$AR** | | | **S462** | |
|---|---|---|---|---|
| | | **$k_1$** | **$k_2$** | **$UU_0$** |
| **S461 [$10^{-3}$/h]** | 1 | 57 | 38 | 67 |
| | 2 | 53 | 36 | 49 |

**Supplementary Table 6. Ranking of normalized curves and used reporter peaks.** Normalization of phosphorylation levels was conducted as previously described by Theillet et al., 2013. (**a**) Reporter peaks for the individual phosphorylation sites are listed. (**b**) Normalized curves were ranked according to the number and quality of peaks which could be included into the calculation of the 100% phosphorylation level:

- **AAA**: Unphosphorylated counterpart and neighboring amino acid were readily available for analysis. Values derived from different reporter peaks matched well (<15% deviation) are marked with (+), values whose values did not match well (>15% deviation) are marked with (-).
- **AA**: Unphosphorylated counterpart or a neighboring amino acid was used for normalization.
- **A**: Only one reporter peak was available, which itself displayed several states or peak overlap.

**a**

| Normalized peaks | Reporter peak |
|---|---|
| pS412 ($\beta_1$AR) | S412, A411, G413 |
| pS459 ($\beta_1$AR) | S459 |
| pS461 ($\beta_1$AR) | S461 |
| pS462 ($\beta_1$AR) | S462 |
| pS473 ($\beta_1$AR) | S473 |
| pS464 ($\beta_2$AR) | S364, Q363 |
| pS345 ($\beta_2$AR) | S345 |

| Normalized peak | Reporter peak |
|---|---|
| pS346 ($\beta_2$AR) | S345 |
| pS343 (Rho) | S343, Q334 |
| pS334 (Rho) | A333 |
| pS338 (Rho) | S338 |
| pT340 (Rho) | E341 |
| pT336 (Rho) | T336 |

**b**

| | AAA | AA | A |
|---|---|---|---|
| **Rho** | pS343(+) | pS334<br>pS338 | pT340<br>pT336 |
| **$\beta_1$AR** | pS412(-) | pS461<br>pS462<br>pS473 | pS459 |
| **$\beta_2$AR** | pS364(+) | pS345 | pS346 |

$$UU \overset{k_1}{\rightarrow} PU \overset{k_2}{\rightarrow} PP$$

*(1.1)*

$$[UU] = [UU_0]e^{-k_1 t}$$

*(1.2)*

$$[PU.1] = \frac{k_1}{k_2 - k_1} * \left(e^{-k_1 t} - e^{-k_2 t}\right) * \ [UU_0]$$

*(1.3)*

$$[PU.2] = \frac{k_1}{k_2 - k_1} * \left(e^{-k_1 t} - e^{-k_2 t}\right) * \ [UU_0] + \left(1 + \frac{k_1 e^{-k_1 t} - k_2 e^{-k_2 t}}{k_2 - k_1}\right) * \ [UU_0]$$

*(1.4)*

$$[PP] = \left(1 + \frac{k_1 e^{-k_1 t} - k_2 e^{-k_2 t}}{k_2 - k_1}\right) * \ [UU_0]$$

*(1.5)*

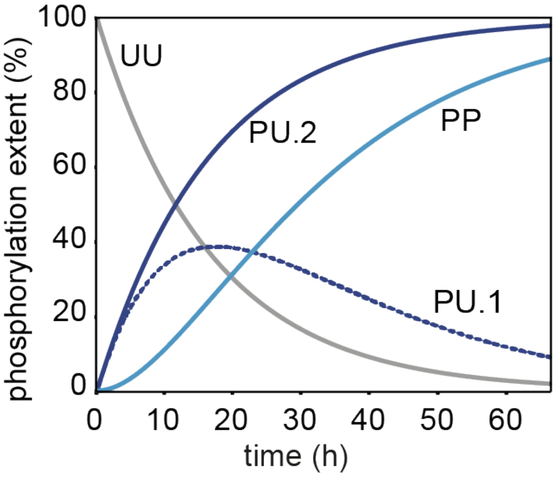

**Supplementary Figure 1. Simulation of relative peak integrals according to formulas 1.1–1.5.** Simulated curves are shown for different states, the unphosphorylated peptide (UU, gray), the mono-phosphorylated peptide (PU.1, dark green and PU.2, light green), and the doubly phosphorylated peptide (PP, purple). PU.1 and PU.2 show the curve progressions for signals, for which the chemical shift is affected or unaffected by the second phosphorylation, respectively. For the simulation the following values were used: $[UU_0]$ =1, $k_1$ = 0.06/h, $k_2$ = 0.054/h.

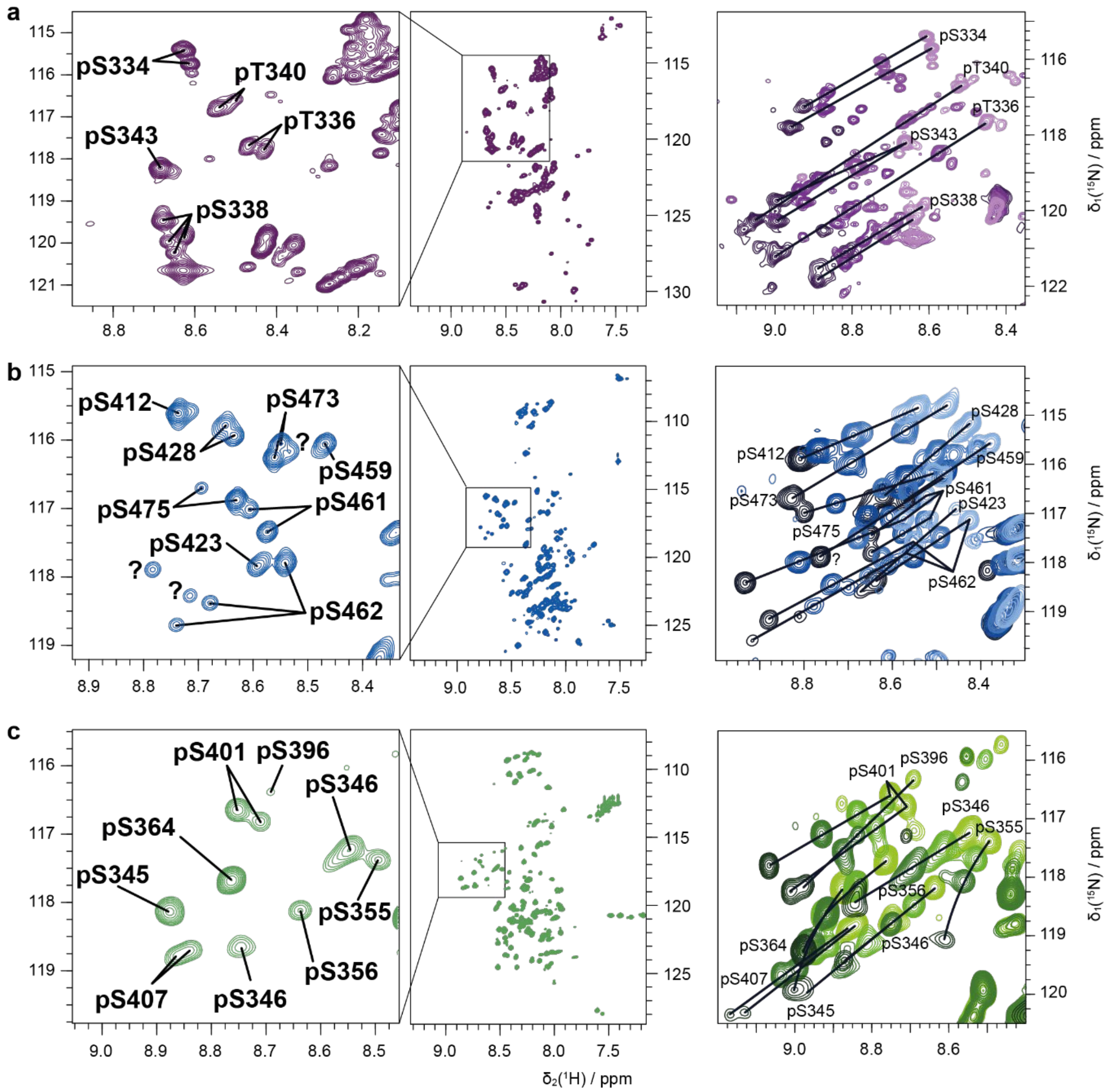

**Supplementary Figure 2. Assignment of the phosphorylated residues of the Rho, $\beta_1$AR and $\beta_2$AR Cterm.** Assignments of the respective phosphorylated amino acid to their peaks are shown for Rho (**a**), $\beta_1$AR (**b**) and $\beta_2$AR Cterm (**c**). On the right-hand side of each panel a pH-titration from 5.5 to 7.4 is depicted (lower pH values are indicated as lighter colors).

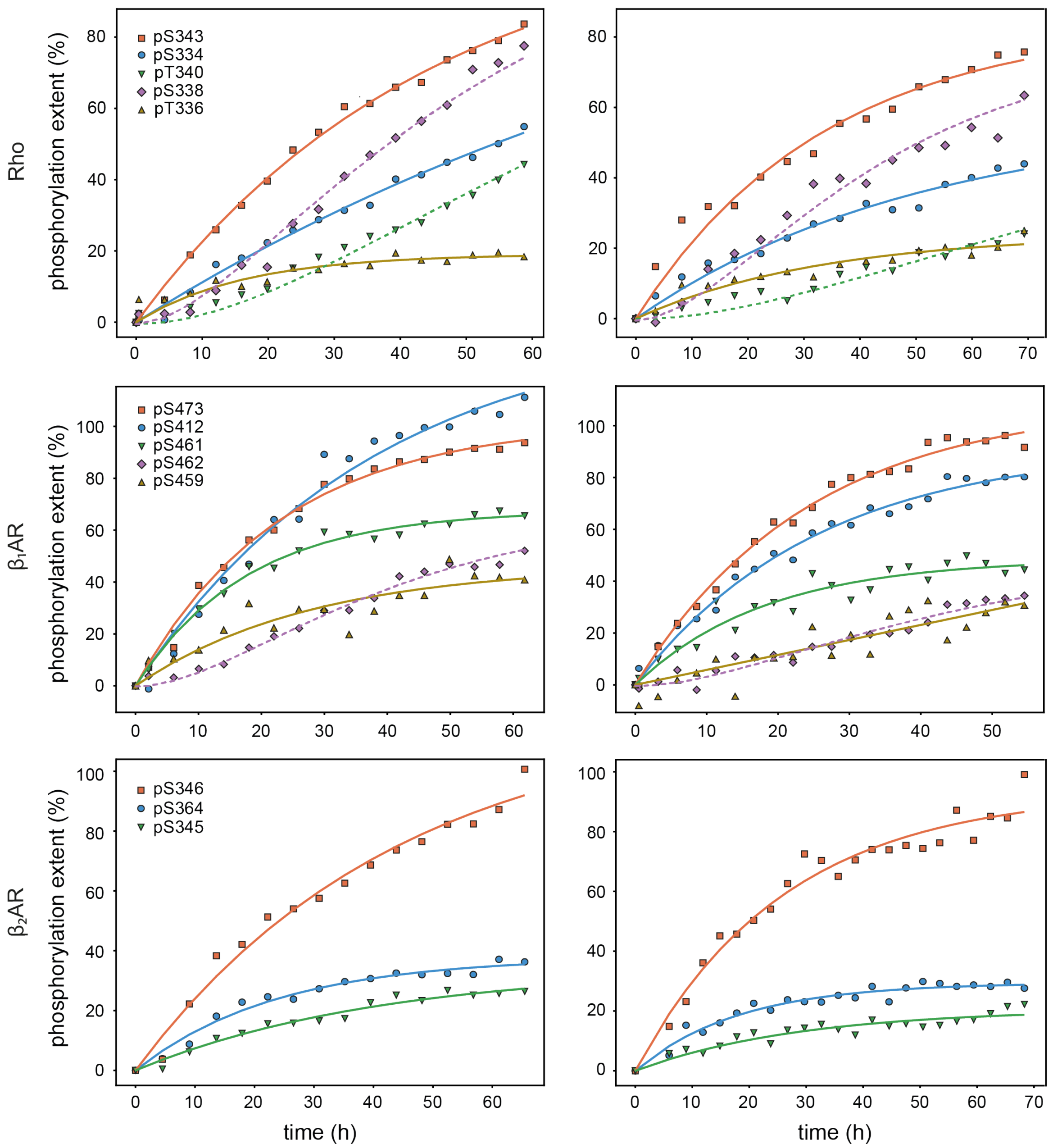

**Supplementary Figure 3. Reproducibility of the phosphorylation reaction of Rho Cterm by GRK1 and β1AR and β2AR Cterm by GRK2.** Duplicate experiments (right column) were measured 1–2.5 years after the initial data sets were recorded (left column). New buffers were prepared, some peptides were freshly purified, a new GRK2 aliquot was prepared, and GRK1 aliquots were used after storage for over 3 years at -80°C. Exponential rising curves are displayed in solid lines. Biphasic curves fitted based on equations 1.1–1.5 are represented with dashed lines. Curves were fitted using JupyterLab4.0.7.

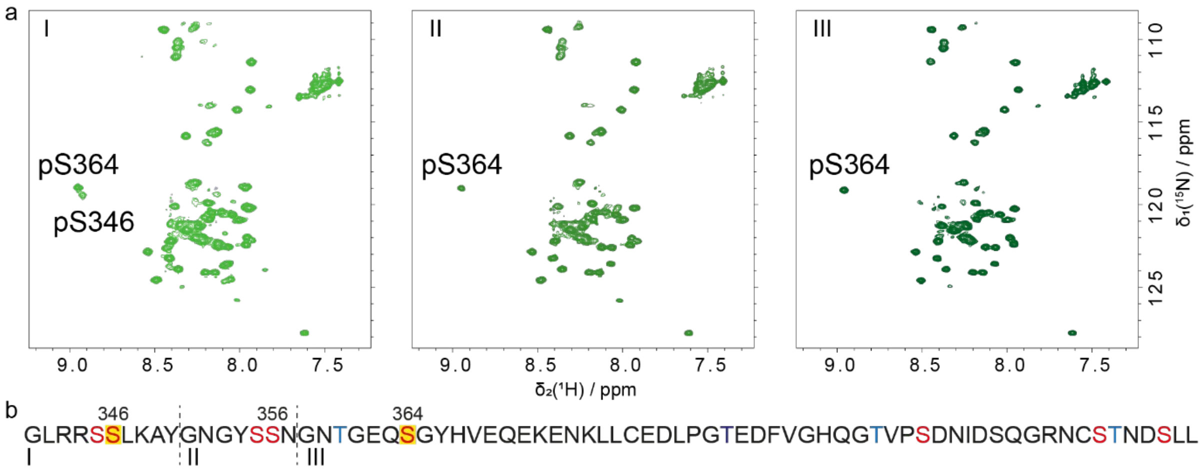

**Supplementary Figure 4. Phosphorylation experiments of three different β2AR constructs confirm site-selectivity of GRK2.** (**a**) 2D [$^{15}$N, $^{1}$H]- HMQC spectra are depicted of three different β2AR C-tail lengths after 30 h of reaction. Used constructs are shown in **b** (I: G342–L413, II: G351–L413, III: G358–L413). Possible phosphorylation sites are marked in red (serines) or blue (threonines). Phosphorylation occurring in this assay are highlighted in yellow.

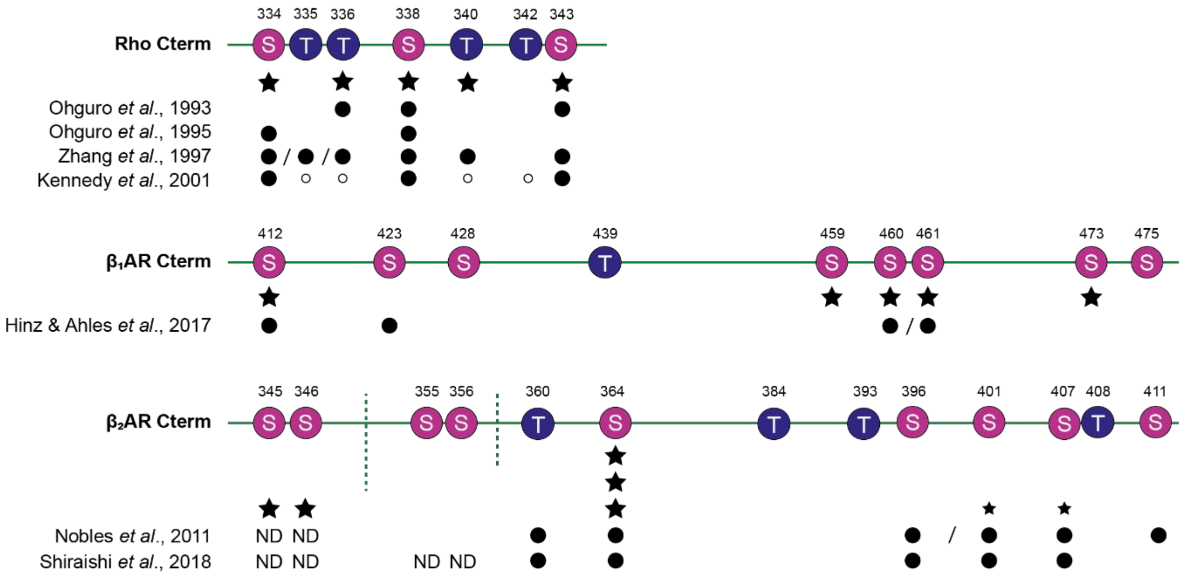

**Supplementary Figure 5. Literature comparison of phosphorylation sites on the Rho Cterm (top), β1AR Cterm (middle) and β2AR Cterm (bottom).** Obtained results within this work are depicted in star shaped forms for the native reaction pairs (Rho:GRK1, β1AR:GRK2, β2AR:GRK2). Stars represent observed phosphorylation within the developed phosphorylation assay. Literature results are marked with filled circles = detected, empty circles = one of the marked residues is phosphorylated, two filled circles separated by a slash = not distinguishable by the applied method, ND = not detectable, no symbol = no observed phosphorylation. Small symbols (stars or circles) represent traces of phosphorylation. Note that the study from Hinz and Ahles *et al*. uses purified receptor from whole cells meaning that the contribution of individual kinases is not dissected.
